# Microglial MyD88-dependent signaling influences extracellular matrix development and interneuron maturation in the hippocampus

**DOI:** 10.64898/2025.12.08.692987

**Authors:** Julia E. Dziabis, Irene O. Jonathan, Benjamin L. Horvath, Grace Zhang, Michael S. Patton, Caroline J. Smith, Dang M. Nguyen, Mahika Jammula, Benjamin A. Devlin, S. K. Monroe, Erica J. Freeman, A. Brayan Campos-Salazar, Madeline J. Clark, Staci D. Bilbo

## Abstract

Parvalbumin interneurons (PVIs) are disrupted across diverse neurodevelopmental disorders, highlighting their vulnerability to developmental perturbations. Inflammation can perturb PVI development and function, and inflammatory mechanisms are often propagated within the brain by microglia. Yet the microglial mechanisms linking inflammatory signals to interneuron development are unclear. To test the role of microglial innate immune signaling in PVI development, we used mice lacking toll-like receptor adaptor MyD88 specifically in microglia. MyD88-deficient microglia showed reduced inflammatory responses but increased early-life phagocytosis of inhibitory synaptic material. In adulthood, males without microglial MyD88 exhibited increased hippocampal PVI density, increased extracellular matrix (ECM) deposition, increased inhibitory signaling, and impaired discrimination behaviors. We determined the cytokine interleukin (IL)-33, which normally drives adult microglial remodeling of the ECM, is developmentally regulated in the hippocampus. MyD88-deficient microglia fail to respond to IL-33, leading to reduced remodeling of the ECM component aggrecan. These results reveal microglial immune signaling via MyD88 regulates hippocampal inhibitory circuit development in a sex-specific manner.

## Introduction

Many neuropsychiatric disorders, while distinct in pathophysiology, symptomology, and onset, share common cellular hallmarks. For example, disruptions to the function of parvalbumin+ interneurons (PVIs), a major subtype of GABAergic neuron, are strongly implicated in schizophrenia, autism spectrum disorder (ASD), Alzheimer’s Disease, and even substance use disorders, such as alcohol use disorder (AUD) (Tang et al., 2021). PVIs are fast-spiking cells that precisely coordinate the firing activity of large networks of neurons to maintain normal brain function. Their extended maturation trajectory and high metabolic needs make PVIs particularly susceptible to perturbations during development, such as immune activation, early life stress, and toxin exposure (Ruden et al., 2021).

Microglia, the resident immune cells of the brain, are also a shared cellular player of interest in the same diverse brain disorders. There is increasing interest in the involvement of immune system dysfunction in these disease mechanisms, as neuroinflammation and immune dysregulation are common features in both animal models and clinical populations. Importantly, during sensitive windows of brain development, microglia interact closely with maturing neurons, influencing their function and survival, and play critical roles in disease-relevant synaptic alterations and later-life behavior (Dziabis and Bilbo, 2022). Only very recently have microglia been shown to be directly involved in postnatal interneuron-specific synaptic and axonal refinement (Favuzzi et al., 2021; Gallo et al., 2022; Gesuita et al., 2022). Additionally, microglia interact with and modulate extracellular matrix (ECM) components via immune signals, with consequences for interneuron function (Wong and Favuzzi, 2023). Therefore, microglia are uniquely poised as integrating nodes of inflammatory signals onto these susceptible neurons and their associated circuits during development.

Understanding how fundamental microglial interactions with interneurons change in response to immune challenge during sensitive developmental windows may provide insight into the etiology of common cellular phenotypes across many diseases. We aimed to broadly modulate microglial pro-inflammatory signaling pathways through cell-specific loss of MyD88, a critical co-adaptor downstream of pattern recognition receptors, such as toll-like receptors (TLRs), and the interleukin-1 family of receptors. We demonstrate that the loss of the MyD88-dependent signaling pathway increases microglial interactions with GABAergic synapses and decreases interactions with the ECM through phagocytic mechanisms in the dentate gyrus of the hippocampus. These mice exhibit long-term changes in PVI synapses, ECM deposition, and GABAergic signaling, with consequences for discrimination behaviors. We implicate neuron-derived cytokine IL-33 signaling through MyD88-dependent pathways in microglia as a key disrupted signal to change developmental microglia interactions with the maturing ECM. Together, we show that microglial inflammatory signaling pathways are critical for hippocampal interneuron development, both in homeostatic and inflammatory conditions, through extracellular matrix interactions.

## Results

### Constitutive loss of MyD88 in microglia blunts acute MyD88-dependent cytokine release, and results in robust transcriptional changes in the absence of inflammatory challenge

Myeloid differentiation adaptor protein 88 (MyD88) is a critical co-adaptor protein downstream of nearly all toll-like receptors (TLRs), as well as the interleukin-1 (IL-1R) family of receptors. MyD88-dependent signaling leads to the activation of transcription factors that prompt the creation and release of cytokines and chemokines to propagate inflammation (Figure 1A). MyD88 is expressed by most cell types in the brain across development, so to manipulate macrophage-specific pro-inflammatory signaling in the early postnatal period, we used a transgenic mouse line to constitutively remove MyD88 from Cx3cr1-expressing cells (Figure 1B). To validate the mouse line, microglia were isolated from male and female CON and cKO mice in the first postnatal week (postnatal day (P)4-6). Isolated microglia were treated with 100ng/mL of lipopolysaccharide (LPS), a potent TLR4 agonist, or sterile PBS (Figure 1C). The TLR4 signaling pathway contains both MyD88-dependent and-independent arms, allowing confirmation of the specificity of MyD88 loss on downstream factor release (Figure 1A). After 4 hours of treatment, LPS increased the MyD88-dependent cytokine TNF-α from CON microglia in supernatant, while cKO microglia treated with LPS did not (Figure 1D). Importantly, LPS led to the release of CXCL10, a MyD88-independent signaling factor, from both CON and cKO microglia (Figure 1E), confirming high specificity of MyD88-dependent signaling removal.

**Figure 1.**
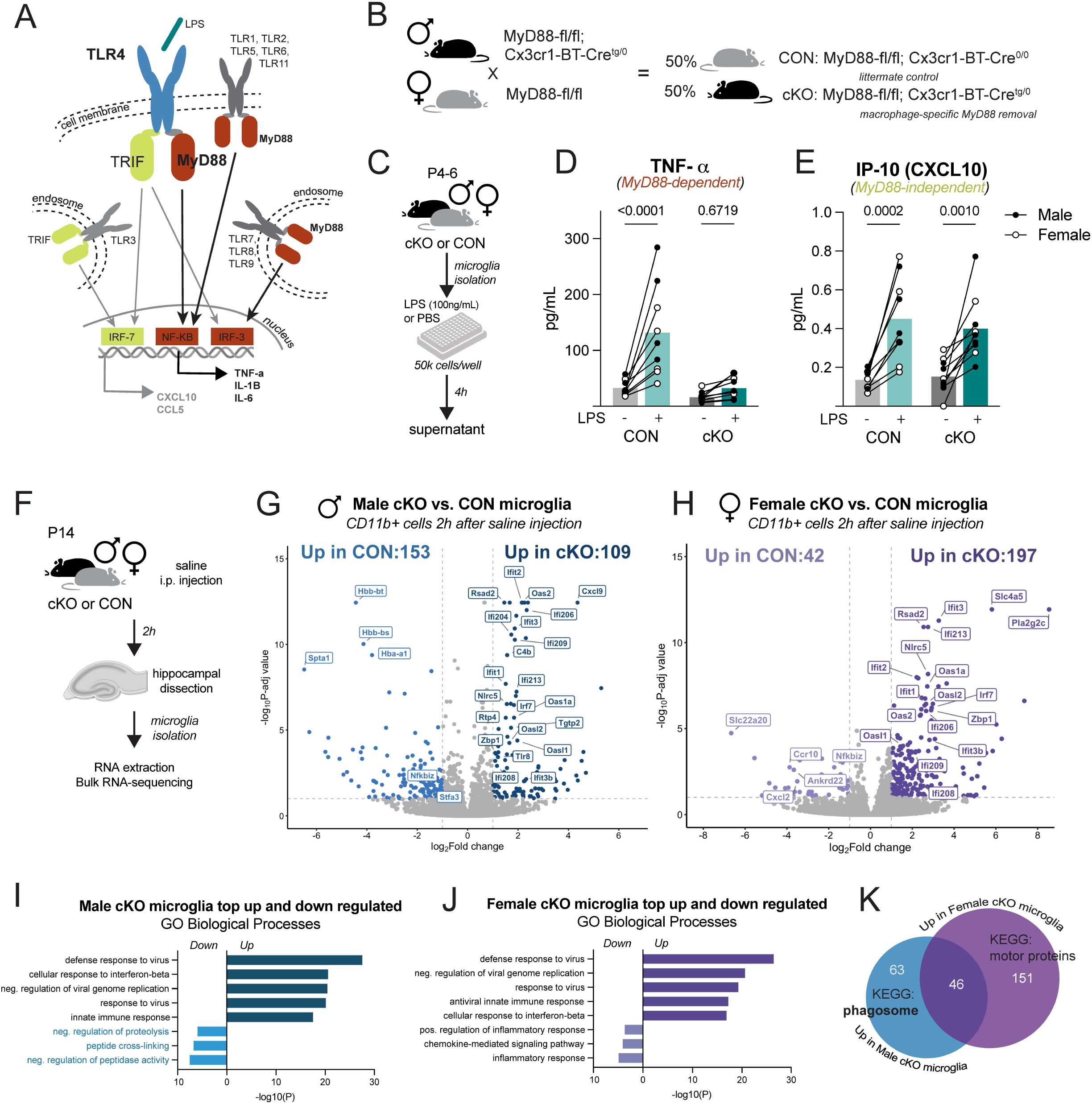
MyD88-cKO microglia release less MyD88-dependent inflammatory factors, and have broadly different gene expression **A.** MyD88 signaling pathway downstream of toll-like receptors (TLRs). **B.** Breeding scheme for constitutive microglial-MyD88 knockout mouse line. **C.** Experimental design for ex-vivo stimulation experiment (LPS = lipopolysaccharide). **D.** TNF-alpha in supernatant 4h after PBS or LPS. 2-way RM ANOVA, Bonferroni post-hoc. **E.** CXCL10 in supernatant 4h after PBS or LPS. 2-way RM ANOVA Bonferroni post-hoc. **F.** Experimental design for bulk RNA-sequencing experiment (i.p. = intraperitoneal injection). **G.-H.** Volcano plot of significantly differentially expressed genes (DEGs) between cKO and CON saline male (**G.**) and female (**H.**) microglia (p-adjusted value cut off of 0.1). I.-J. Gene ontology analysis of top differentially expressed genes in male (**I.**) and female (**J.**) cKO microglia. **K.** Overlapping and unique upregulated genes in cKO males and females, and top unique gene KEGG pathways.

We next determined if loss of MyD88 in microglia altered their gene expression profiles. In brain regions such as the hippocampus, normal microglial developmental interactions with neurons like synaptic refinement peak around the end of the second postnatal week. To assess transcription at this developmental time point, microglia from the hippocampi of male and female CON and cKO mice were isolated at P14 for bulk RNA-sequencing (Figure 1F). In males, differential gene expression analysis revealed 153 genes that were significantly downregulated in cKO microglia compared to CON, and 109 genes that were upregulated by cKO microglia compared to CON (Figure 1G). Similarly, 42 genes were downregulated by cKO female microglia, and 197 genes were upregulated in cKO female microglia compared to CON (Figure 1H). Using gene ontology analysis to highlight major gene expression patterns, we identified many common upregulated biological processes in both male and female cKO microglia related to viral responses (Figure 1I-J). We hypothesize this strong anti-viral phenotype seen across both sexes to be compensation through MyD88-independent signaling pathways in cKO microglia, such as via TRIF (Figure 1A). In females, top downregulated biological processes were all related to inflammation and chemokine-mediated signaling (Figure 1J), confirming the effects of the loss of MyD88-dependent signaling. However, in male cKO microglia, the top downregulated biological processes were all related to the stability, processing, and degradation of proteins (Figure 1I). These findings suggest that male cKO microglia increase their proteolytic activity (by downregulating inhibitors of proteolysis and peptidases), leading us to hypothesize that in cKO males, there may be increased capacity for remodeling and clearance of materials such as synapses, protein aggregates, or extracellular matrix (ECM) components. The additional “peptide cross-linking” function downregulation further supports the idea that cKO male microglia may be highly involved in ECM degradation and synapse destabilization compared to CONs. When comparing gene expression changes between the sexes, nearly 50 of the significantly upregulated DEGs were shared by male and female cKO microglia (Figure 1K). When the male-cKO unique genes were run through KEGG pathway analysis, the top hit was “phagosome,” compared to “motor proteins” in females (Figure 1K). Together, these data suggest that loss of MyD88 from microglia affects inflammatory gene expression in both sexes, but male microglia appear to be uniquely impacted in functions related to phagocytic and proteolytic activity.

### Male microglia lacking MyD88 exhibit altered phagocytosis during the second postnatal week

With the unique proteolytic and phagocytic gene expression changes in mind, we next determined if male cKO microglia differ functionally from CON microglia across the developmental synaptic refinement window in the hippocampus (Figure 2A). We initially chose to focus on microglia from the dentate gyrus, as this region contains a high density of diffuse ECM, and microglia are known to influence ECM deposition in the adult dentate gyrus at least partially through phagocytic mechanisms (Bolós et al., 2018; Nguyen et al., 2020). We hypothesized that male cKO microglia would be more phagocytic than controls around the second postnatal week, based on previous developmental microglial synaptic interaction work in the hippocampus that focused on P14-17 as the peak of synaptic interactions in CA1 (Dayananda et al., 2023; Paolicelli et al., 2011; Weinhard et al., 2018). Individual microglia from the P12, P15, and P18 dentate gyrus were volumetrically reconstructed to determine the percent of their total volume filled by CD68+ lysosomes (Figure 2B). Microglia from cKO males contained a greater percent of CD68+ volume than CONs at P12 and P15 (Figure 2C-D), with CON lysosome volume the greatest at P12. By P18, genotype differences in lysosomal volume resolved (Figure 2E).

**Figure 2.**
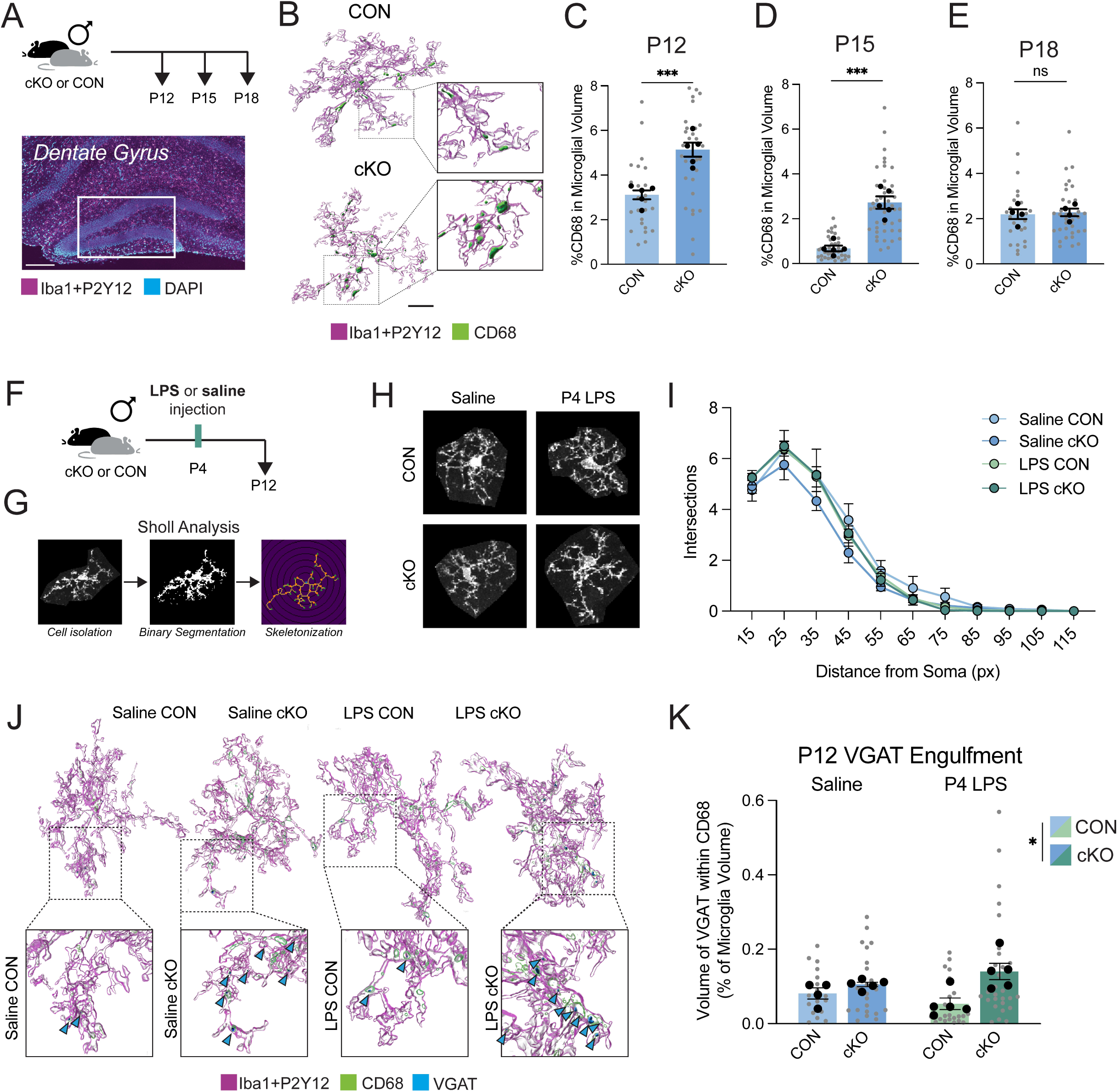
MyD88-deficient microglia preferentially engulf inhibitory synaptic material at postnatal day 12 **A.** Experimental design for microglial phagocytic capacity experiment and region of interest within the dentate gyrus. Scale bar = 200μm. **B.** 3D reconstructed examples of CD68 (lysosome) in P12 CON and cKO microglia. Scale bar = 10μm. **C.-E.** Quantification of lysosomal content within dentate gyrus microglia at P12 (**C.**), P15 (**D.**), and P18 (**E.**). Asterisks indicate significant by unpaired t-test, n = 4-5 animals/group. **F.** Experimental design for P12 microglial endpoints. 330mg/kg dose of LPS was administered at postnatal day (P)4. **G.** Method overview for Sholl analysis endpoints. **H.** Representative individual microglia from Sholl analysis. **I.** Quantification of process complexity by Sholl analysis. Non-significant by repeated measures 3-way ANOVA, n = 4-5 animals/group. **J.** Representative images of microglia engulfment of VGAT synaptic material, indicated by blue arrows. **K.** VGAT engulfment, significant main effect of genotype by 2-way ANOVA, n = 4-5 animals/group. All error bars = SEM, all statistics run on individual animals (black or black-outlined circles), grey points represent individual microglia analyzed.

As the age with the greatest lysosomal volume in both CON and cKO microglia, we next wanted to further characterize P12 dentate gyrus microglia across genotypes. To potentially “unmask” any effects of the loss of MyD88 signaling and to model the impact of early-life inflammation, we introduced a group of CON and cKO mice treated with a moderate dose of LPS (330μg/kg) or sterile saline subcutaneously at P4 before collecting brains at P12 (Figure 2F). LPS is a known activator of TLR4-dependent signaling, the response to which we confirmed is blunted in cKO microglia in culture (Figure 1D). We first found that neither P4 LPS nor loss of MyD88 resulted in broad changes in dentate gyrus male microglia process complexity at P12 by Sholl analysis (Figure 2G-I). Further, the density of male microglia across the dorsal hippocampus was not changed across conditions at P12 (Figure S1A), even though microglia numbers are likely still expanding at this age (Kim et al., 2015). To quantify synaptic engulfment, we volumetrically reconstructed microglia, measuring the amount of presynaptic material within their lysosomes, as postsynaptic material is less likely to be engulfed at this developmental age (Weinhard et al., 2018). First, knowing that VGlut2+ synaptic material is engulfed by microglia in other brain regions in the early postnatal window (Block et al., 2022; Devlin et al., 2025), and that VGlut2+ synapses are reduced in the hippocampus as VGlut1+ synapses are increased (Miyazaki et al., 2003), we quantified VGlut2+ inclusions in microglial lysosomes. Very little VGlut2+ material was observed in male microglia lysosomes at P12 in the dentate gyrus (Figure S1B). However, microglia are also critical remodelers of inhibitory synapses and axons from PVIs and somatostatin (SST) expressing interneurons around this developmental window in other brain regions, e.g. somatosensory cortex (Favuzzi et al., 2021; Gesuita et al., 2022). In the P12 dentate gyrus, cKO microglial lysosomes contained more vesicular GABA transporter (VGAT+), which is used to broadly identify inhibitory presynaptic elements, with the greatest inclusion observed in mice without MyD88 that received LPS challenge at P4 (Figure 2J-K). Interestingly, P12 female MyD88-cKO microglia also contained significantly more VGAT material within their CD68+ lysosomes, suggesting that loss of MyD88 similarly impacts male and female microglial phagocytic changes at P12 (Figure S1C). These effects were not driven by any changes in the overall volume of cells in either male or female microglia (Figure S1D), or by increases in the expression of functional GABA_B_ receptors on cKO microglia, as has been previously described in the barrel cortex and measured by expression of both *Gabbr1* and *Gabbr2* within microglia (Favuzzi et al., 2021)(Figure S1E-F). Together, these data suggest that microglia lacking MyD88 have greater lysosomal volume and engulfment of inhibitory presynaptic material during critical windows of synaptic refinement.

### PVI-specific inhibitory synapses and inhibitory signaling are increased in the MyD88-cKO male hippocampus

We next determined if the increase in VGAT+ synaptic material within MyD88-deficient cKO microglia observed at P12 led to changes in inhibitory synapse density in the dentate gyrus. We hypothesized that increased VGAT engulfment at P12 would result in reduced inhibitory synaptic density at P18. Using the same P4 LPS paradigm (Figure 3A), we labeled tissue with synaptic protein pairs and the colocalization of pre-and post-synaptic proteins was quantified as a proxy for complete synapses (Figure 3B). We first quantified VGAT+/gephyrin+ synapses in the P18 male dentate gyrus and found no changes in inhibitory synapses (Figure S1G). However, VGAT labels all GABAergic synapses, and in the somatosensory cortex, there are known differences in microglial interactions with different subtypes of GABAergic neurons (Favuzzi et al., 2021; Gesuita et al., 2022). Using Synaptotagmin-2 (Syt2) as a PVI-specific presynaptic marker when colocalized with gephyrin (Sommeijer and Levelt, 2012; Favuzzi et al., 2021; Irala et al., 2024), we found that MyD88-cKO mice had more PVI synapses at P18 than CONs (Figure 3C-D). Despite increased VGAT engulfment in both male and female cKO microglia (Figure 2K, Figure S1C), the increase in Syt2+/gephyrin+ synapses in cKO mice at P18 was specific to male mice, as we did not see any differences in synaptic density in P18 females (Figure S1H). There were also no changes in male VGlut2+/PSD95 excitatory synapses at P18, which was expected due to the lack of changes in VGlut2+ synaptic internalization in lysosomes at P12 (Figure S1I).

**Figure 3.**
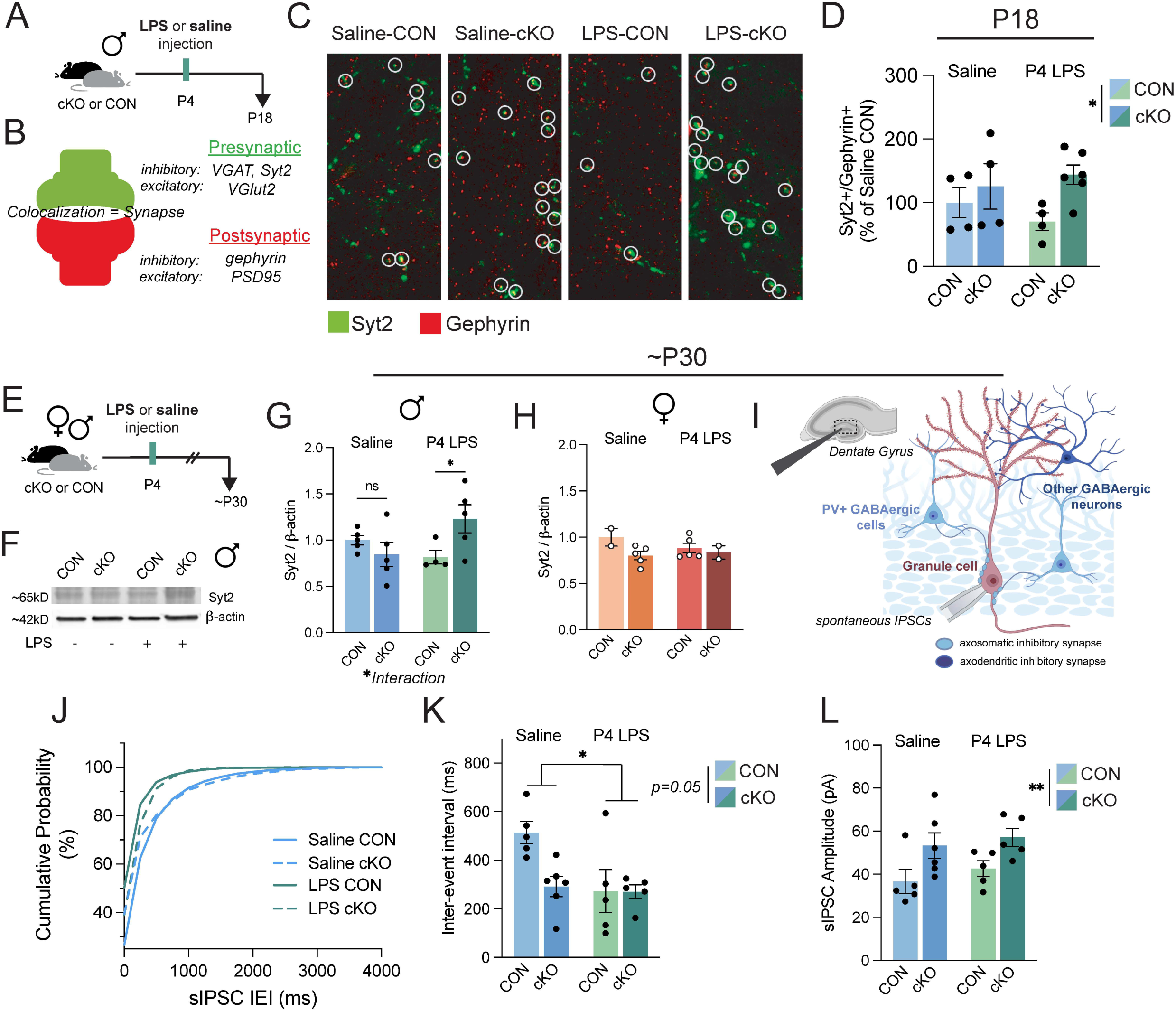
Hippocampal dentate gyrus inhibitory signaling is altered in males without microglial MyD88 **A.** Experimental design for synapse density quantification at P18 following P4 endotoxin injection. **B.** Representative schematic of pre-and post-synaptic protein colocalizations. **C.** Representative images of PV-specific synaptic density, circles indicate colocalized pre-(Syt2) and post-(gephyrin) synaptic proteins. **D.** Quantification of Syt2+/gephyrin+ synaptic density, normalized as percent of Saline-CON, significant main effect of genotype by 2-way ANOVA, n = 4-6 animals/group. **E.** Experimental design for ∼P30 collections for protein abundance by western blot and electrophysiology endpoints. **F.** Representative blot of Syt2 protein from male hippocampi. **G.-H.** Quantification of Syt2 protein normalized to β-actin abundance in males and females. Non-significant changes in females (**H.**), main interaction effect by 2-way ANOVA in males (**G.**), asterisk indicates significant Bonferroni post-hoc, n = 2-5 mice/group. **I.** Representative schematic of electrophysiology from dentate gyrus slice to quantify spontaneous inhibitory post-synaptic currents (sIPSCs) onto granule cells in males. **J.** Cumulative probability of inter-event intervals (IEI), significant by Kruskal-Wallis test. **K.** Quantification of IEI (frequency), significant main effect of treatment by 2-way ANOVA, trending main effects of genotype (p=0.05) and genotype x treatment interaction (p=0.05). **L.** Amplitude of sIPSCs, significant main effect of genotype by 2-way ANOVA. n = 5-6 mice/group, 1 point = average of 1-3 cells/animal. All error bars = SEM, all statistics run on individual animals.

One alternative hypothesis that emerged in the wake of the unexpected abundance of Syt2+ synapses was that the increased VGAT engulfment seen in both male and female cKO microglia was indicative of a phagocytic deficit, rather than an increased clearance of material. Microglia under stress can engulf, but fail to break down proteins in their lysosomes, which may at a single time window snapshot appear to be active remodeling, when in fact we are capturing debris build-up (Quick et al., 2023). To test this hypothesis, we isolated microglia from CON and cKO P12 male and female mice, allowed the cells to adhere in culture, then imaged their inclusions of pHrodo *E. coli* over time (Figure S2A). PHrodo will emit fluorescence when included in a low pH environment, such as a microglial lysosome, as microglia in culture uptake and break down the attached *E. coli*. We found that cKO male microglia are just as efficient at taking up *E. coli* into their lysosomes, and non-significantly more efficient at breaking down material, compared to CON microglia (Figure S2B). In contrast, cKO female microglia were slightly less efficient at acute uptake but broke down material at a similar level (Figure S2D). We next compared the breakdown of more “immune-neutral” material in late endosomes at the timepoint of peak uptake, which was around 3 hours after treatment for both males and females. We applied DQ-BSA to adhered microglia in culture (Figure S2A). In this assay, fluorescence is observed only when late endosome proteolysis of dye-labeled BSA peptide occurs. We quantified the percent of total microglia (quantified by NucBlue Hoescht stain) that contained fluorescence at 3 hours after treatment and found more cKO male microglia were fluorescent than CONs (Figure S2C). In contrast, female cKO microglia were less likely to be fluorescent compared to CONs (Figure S2E). Taken together these data suggest male cKO microglia are if anything *more* efficient at phagocytosis and lead us to believe that the inclusions of VGAT+ synaptic material in male cKO lysosomes were indeed being actively broken down, and not due to a deficit in endosomal breakdown. These culture experiments also reveal a possible explanation for the sex differences observed later in life; while there was increased VGAT and lysosomal content within female cKO microglia compared to controls, our findings suggest that female cKO microglia are not as efficient as female CONs at uptake and breakdown of material. This snapshot of increased synaptic engulfment at P12 in females may therefore be compensatory for these microglia to “catch up” to CON levels of degradation and may be the reason behind the normalized synaptic density between genotypes in females by P18.

Importantly, microglia can impact inhibitory synaptic development by more than just synaptic removal; in the developing somatosensory cortex, microglia regulate axo-axonic synaptogenesis (Gallo et al., 2022). The authors suspected that this is mediated by microglial-derived synaptogenic factors, though microglial impacts on PVI synapses specifically occur only after the initial assembly of these synapses, after P12 (Favuzzi et al., 2021). While there were no sequencing differences in MyD88-cKO microglia that point to changes in synaptogenic factors, we wanted to confirm that MyD88 loss in microglia did not lead to changes in inhibitory synapse density prior than P18, which could influence microglial engulfment of VGAT+ synaptic material. Quantification of both VGAT+/gephyrin+ synapses and PVI-specific Syt2+/gephyrin+ synapses in males at P12 revealed no significant differences between CON and cKOs, suggesting that the increase in PVI synapses emerges after P12 (Figure S2F-G).

At P18, both the GABAergic system and the hippocampus are still developing. PV protein is only just being expressed, extracellular matrix structures are still being constructed and refined, and precise adult memory engrams do not mature until P24 in mice (Pelkey et al., 2017; Ramsaran et al., 2023). Around 1 month of age, the hippocampal inhibitory system is thought to be functionally mature, so we again followed the same experimental paradigm (Figure 3E) and collected tissue from brains of 1 month old mice for either western blot protein quantification (Figure 3F) or electrophysiology (Figure 3I). We first determined that P30 cKO males that received P4 LPS continued to have elevated levels of Syt2 protein (Figure 3G). We again confirmed these inhibitory changes were male-specific, as female cKO mice did not have more Syt2 protein at P30, similar to P18 findings (Figure 3H). We next determined if the increase in Syt2 inhibitory synapses in cKO males resulted in long-term functional consequences in hippocampal signaling by electrophysiology. We hypothesized that this increase in presynaptic protein, if indicative of increased synaptic function, would result in more frequent inhibitory events in the male dentate gyrus. Using pharmacology to block excitatory currents, we isolated and recorded spontaneous inhibitory post-synaptic currents (sIPSCs) by whole-cell patch-clamp recording from granule cells in the dentate gyrus (Figure 3I). We found that the frequency of sIPSCs was increased in P4 LPS treated animals, though there was a trending (p = 0.05) main effect of genotype and interaction effect (p = 0.05) (Figure 3J-K). Additionally, the amplitude of sIPSCs was greater in the cKO dentate gyrus than in CONs (Figure 3L). With changes in both amplitude and frequency of sIPSCs, we suspect there is likely a combination of both pre-and post-synaptic changes in GABAergic transmission, such as through an increased number of active GABAergic synapses, rather than solely pre-synaptic Syt2 observed by western blot (Figure 3G).

Additional explanations for these changes could be impacts on GABA_A_ receptor sensitivity or altered chloride homeostasis from a delayed or immature GABA switch. To investigate, we performed additional western blot quantification of ion transporters NKCC1 and KCC2 from the P30 hippocampus (Figure S3A-B). During the early postnatal GABA switch, where GABA becomes hyperpolarizing, the amount of KCC2 dramatically increases and NKCC1 abundance drops. We found a non-significant but trending (p = 0.08) increase in the NKCC1:KCC2 ratio in cKO males, likely driven by the reduced KCC2 abundance in cKO males (Figure S3C-E). These effects were not observed in females (Figure S3F-H). This might point to GABA as less hyperpolarizing (or even depolarizing) in P30 male cKO mice, leading to increased neuronal excitability that may drive a compensatory upregulation of Syt2+ synapses to balance the system. The trending difference also suggests delayed maturation and may explain the electrophysiological findings at the same age: as the ratio recovers to mature levels, extra Syt2+ synapses would produce very strong inhibition.

### Adult dentate gyrus PV interneurons are impacted in MyD88-deficient male mice

Given the increases in PVI synaptic proteins and overall inhibitory signal at 1 month of age in male cKO mice, we next determined if there were PVI changes later in life. In the dentate gyrus, fast-spiking PVIs are critical for complex cognition like learning and memory. Mature PVI somas are often surrounded by perineuronal nets (PNNs), which are specifically organized, non-diffuse ECM structures that protect PVIs, regulate their activity, and impact synaptic plasticity and stabilization (Fawcett et al., 2019). In the visual cortex, for example, PNNs are well known for their critical role in the closure of the critical period of ocular dominance plasticity, and manipulation through degradation of the PNNs can re-open this plasticity (Hensch, 2005; Pizzorusso et al., 2002). Therefore, the increased inhibitory-specific synaptic interactions (Figure 2K), paired with the ECM-remodeling gene expression changes seen in male cKO microglia (Figure 1I), led us to hypothesize that MyD88-cKO mice may have long-term changes to the PVIs and PNNs. Therefore, we injected male CON and cKO mice with LPS or saline at P4 and collected their brains at 3 months old (∼P100) to quantify PV protein expression as well as the expression of PNNs by an antibody against wisteria floribunda albumin (WFA) (Figure 4A-B). We observed an increase in the density of PVIs in the dentate gyrus in adult cKO mice, with the greatest density observed in the P4 LPS-cKO group, consistent with our findings at P30 (Figure 4C-D). We also quantified the density of SST interneurons in the dentate gyrus, as SST cells are born from the same embryonic origin as PVIs and are another major GABAergic cell type in the hippocampus. The density of SST+ cells in the dentate gyrus is very low but unchanged across genotype and treatment, suggesting this is a PVI-specific effect (Figure S3I). If we quantify the ratio of PV cells to SST cells across the whole dorsal hippocampus, we again see a main effect of genotype on the ratio of PVIs:SST cells, where cKO males have a greater amount of PVIs (Figure 4E).

**Figure 4.**
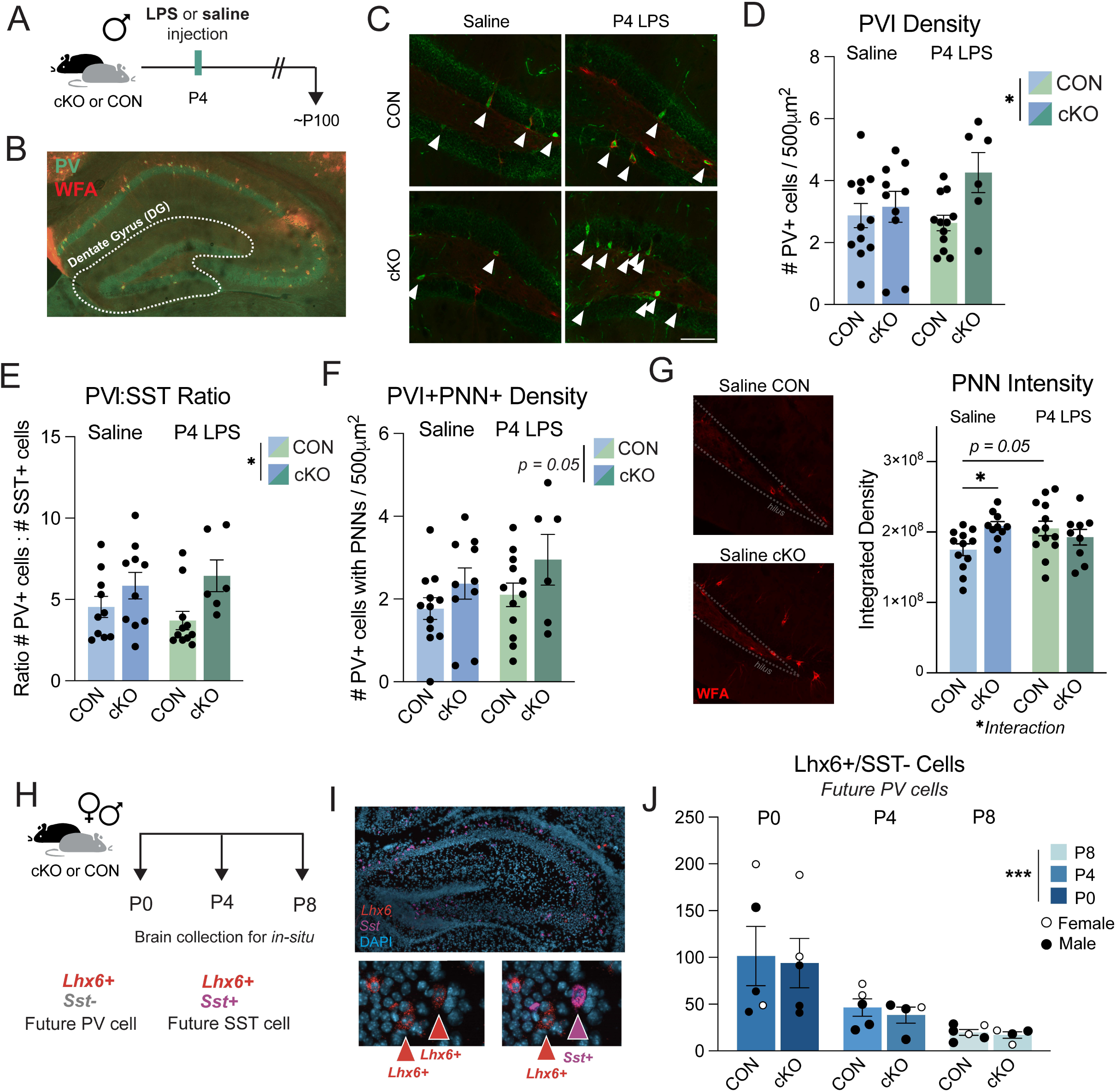
Adult MyD88-cKO males have more hippocampal PVIs than CONs with greater perineuronal net deposition **A.** Experimental design for adult cell quantification in the hippocampus following P4 LPS in males. **B.** Representative image of parvalbumin (PV) interneurons and WFA (perineuronal nets, PNNs) in the dorsal hippocampus, white outline represents area quantified for dentate gyrus (DG) endpoints. **C.** Representative images of PVIs and PNNs in the DG, white arrows represent PV cells. Scale bar = 100μm **D.** Density of PV+ cells across the DG normalized to 500μm^2^ area, significant main effect of genotype by 2-way ANOVA. **E.** Ratio of PVIs to somatostatin (SST) interneurons across the dorsal hippocampus, significant main effect of genotype by 2-way ANOVA. **F.** Density of PVI+/PNN+ cells in the DG normalized to 500μm^2^ area, trending (p=0.05) main effect of genotype by 2-way ANOVA. **G.** WFA labeling for PNNs in the dentate gyrus and quantification of intensity of WFA fluorescence, significant interaction by 2-way ANOVA, asterisks indicate significant Bonferroni post-hoc. For all adult experiments, 1 point = 1 animal, n = 6-12 mice/group, averaged from at least 2 images/animal. **H.** Experimental design for fluorescent in-situ experiment from P0-P8. *Lhx6*+/*Sst*-/DAPI+ cells were quantified as putative future PVIs, *Lhx6*+/*Sst*+/DAPI+ were quantified as putative future SST+ interneurons. **I.** Representative image of P8 dorsal hippocampus, left inset representing *Lhx6*+ cells (red arrows), right inset representing *Sst*-(red arrow) and *Sst*+ cells (magenta arrow). **J.** Quantification of future PVIs across P0-P8, significant main effect of age by 2-way ANOVA. n = 4-5 mice/age/genotype. All error bars = SEM, all statistics run on individual animals.

The colocalization of PNNs, organized peri-somatic ECM structures, with PVIs in adulthood is often used as a measure of mature, functional PVIs. We next determined if there was a change in this subtype of ECM in later adulthood. We observed a trending (p = 0.05) increase in the density of PVIs with PNNs surrounding them (Figure 4F), but no differences in the total density of PNNs in the dentate gyrus (Figure S3J). This lack of change in the *number* of PNNs was not entirely surprising, as many have previously reported structural or compositional changes in PNNs in response to challenge without impacting the overall density of nets. Therefore, we next quantified the intensity of WFA signal from the same sections and found a statistically significant interaction between genotype and treatment, where both genotype and treatment moderately increase the intensity of PNNs (Figure 4G). Together, these data illustrate an adult system with a higher density of PVIs with increased deposition of PNNs around them.

One possible explanation for the increase in PVI density in adulthood is a genotype-induced deficit in the development of hippocampal PVIs. PVIs and SST interneurons are born in the medial ganglionic eminence (MGE) embryonically, then undergo a protracted tangential migration through the cortex and eventually into their final home in the hippocampus. Gestational manipulations directly to microglia or through maternal immune activation impact the migration and eventual laminar distribution of interneurons in the somatosensory cortex (Squarzoni et al., 2014; Thion et al., 2019). Additionally, interneurons undergo a period of cell death in the early postnatal window (Southwell et al., 2012; Wong et al., 2018), and microglia are important for developmental apoptosis and cellular clean up specifically in the dentate gyrus (Cunningham et al., 2013; Sierra et al., 2010; Wakselman et al., 2008). PV protein is not expressed until the second postnatal week, though it is thought that interneuron cellular fate is determined prior to PV expression. Therefore, we designed an experiment to quantify “future” PV-expressing cells in the first postnatal week, when interneuron density is known to drastically drop (P0-P8) (Figure 4H). Using fluorescent in-situ RNAscope, we used the expression of *Lhx6*, which labels all MGE-derived interneurons (primarily PVIs and SST interneurons), and an *Sst* probe to identify the population of *Lhx6*+/*Sst*-cells, which will become PV+ later in life (Figure 4I). We found no differences in the density of “Future PVIs” at P0, confirming there were no broad deficits in interneuron migration to the hippocampus (Figure 4J). We also observed the expected reduction in density of cells across developmental age in both CON and cKO animals, supporting the hypothesis that the changes observed later in life are not due to changes during the cell death period (Figure 4J). There were also no genotype differences in *Lhx6*+/*Sst*+ cells (“Future SST+ cells”) across development (Figure S3K). These results suggest that changes in the density of PVIs in our MyD88-cKO model are not occurring embryonically or in the first postnatal week, but rather later in PVI development.

### MyD88 deficiency in microglia protects females, but not males, from behavior changes following early life immune challenge

We next determined if MyD88 loss in microglia was sufficient to induce behavioral changes in adulthood. Manipulations that impact microglial interactions with neurons during critical developmental windows can have long-term effects on behavior, in both classic somatosensory circuits and associated behaviors, as well as more complex behaviors, such as social behavior (Andoh and Koyama, 2021; Ferro et al., 2021). Therefore, we performed behavioral assays to assess the impact of both early life inflammation and microglial-MyD88 deficiency on male and female behavior (Figure 5A). We hypothesized that male cKO mice may have deficits in dentate gyrus-related behaviors, such as those that require discrimination of similar contexts. We found no changes to locomotor behavior, anxiety-like behaviors, or working memory in any mice (Figure S4A-C). Additionally, when we probed social behaviors, we saw no changes in male or female sociability (preference for a novel conspecific vs. an object) in the 3-chamber social assay (Figure S4D). However, when we tested the social novelty preference of these mice, we found a significant effect of LPS in males, where LPS treatment significantly reduced the preference for the novel mouse, an effect mostly driven by the cKO-LPS group (Figure 5B). To determine if this lack of preference for the novel social stimulus in the male LPS-cKO mice was due to specific changes in social or reward behavior, or a broader deficit in memory or decision-making, we next used the novel object location (NOL) paradigm. In contrast to novel object recognition, which is not dependent on the hippocampus (Barker and Warburton, 2011), NOL uses two of the same objects and instead tests spatial memory through an inter-trial interval between the initial investigation and moving one object to a new location. Discrimination between two similar environments is also a highly dentate gyrus dependent behavior (van Dijk and Fenton, 2018). We again saw no significant effects of treatment or genotype on female behavior, but LPS significantly reduced NOL preference in male mice, with LPS-cKO males having the lowest average performance (Figure 5C).

**Figure 5.**
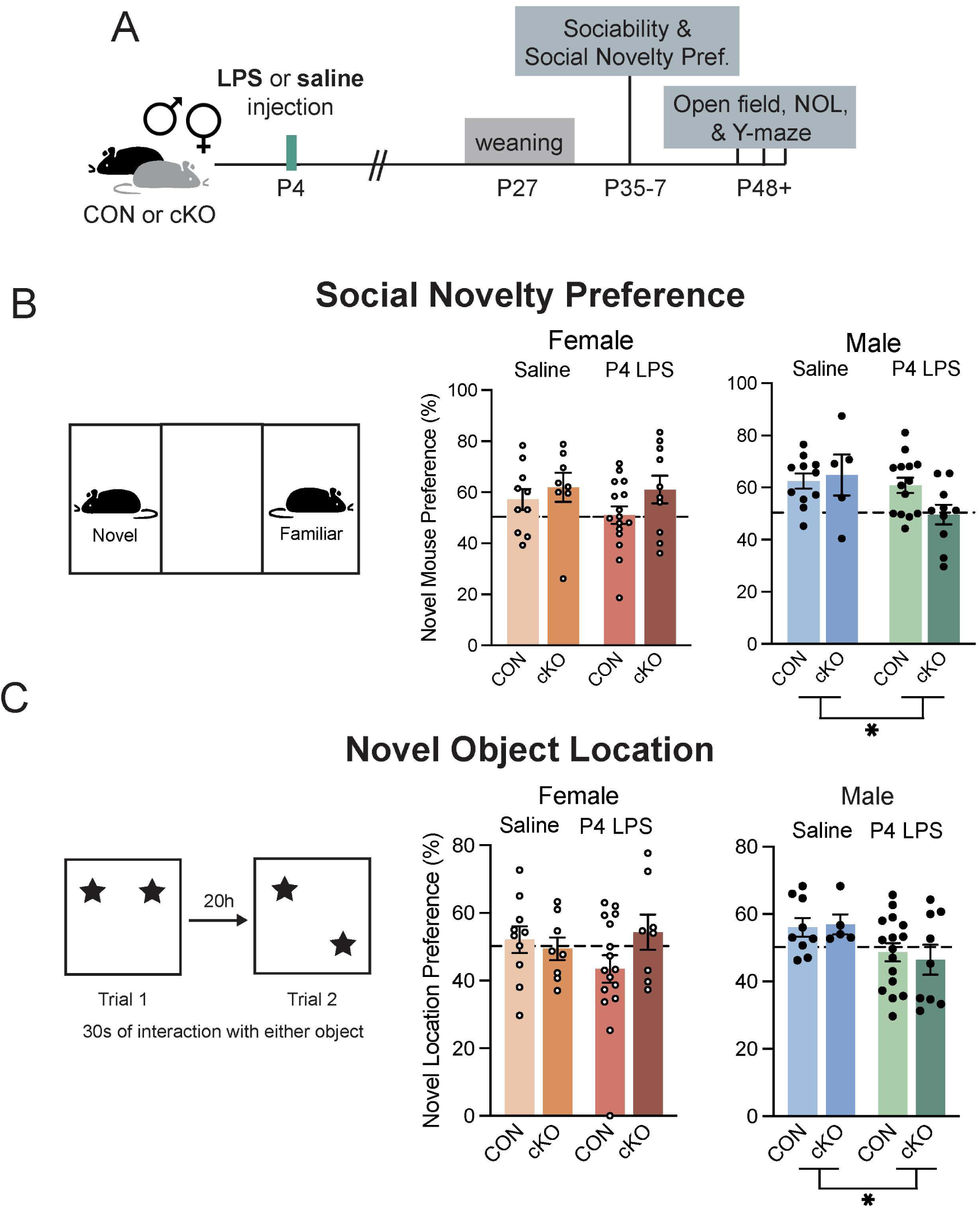
Early life LPS injected males without MyD88 exhibits the greatest discrimination behavior deficits, in both social and non-social behavior paradigms **A.** Experimental design for behavioral characterization experiments with P4 LPS. **B.** Social novelty preference, run immediately following sociability assay (Supp. Fig. 4). Preference for novel mouse determined by percent of time spent investigating the novel mouse over total time spent investigating either mouse. No significant changes in female mice, main effect of treatment by 2-way ANOVA in male mice. **C.** Novel object location task, preference for novel location determined by percent of time spent interacting with the novel location object over total interaction time of the first 30 seconds of interactions with either object. No significant changes in female mice, main effect of treatment by 2-way ANOVA in males. n = 5-16 mice/group. All error bars = SEM, all statistics run on individual animals.

Due to the results of our sequencing experiment suggesting a potential compensatory increase in MyD88-independent signaling pathways (Figure 1G-J), we next tested the hypothesis that the adult changes in male brains and behavior may be due to an increase in these secreted signals (e.g., interferons). We ran a cohort of mice where instead of inducing inflammatory challenge using P4 LPS injection in MyD88-CON and-cKO mice, we injected wild-type mice with polyinosinic:polycytidylic acid (poly I:C) daily from P2-P6 (as in (Schwabenland et al., 2023)). Poly I:C is a synthetic double-stranded RNA that activates TLR3, one of the only toll-like receptors that is entirely MyD88-independent and leads to an increase in type I interferon expression, the family of signals that are upregulated by our cKO microglia (Figure 1A, G-J). We found, however, that there were no changes in social discrimination behavior in early life poly I:C injected males and females, as well as no changes in adult PVI density in these mice (Figure S3E-K). This led us to believe that the observed changes in P4 LPS-treated cKO males were not due to the increased interferon expression in the early postnatal period. Together, these behavioral assessments point to male-specific long-term changes in discrimination-dependent behaviors (Figure 5B-C). This may have relevance to understanding why and how neurodevelopmental disorders with immune contributions tend to be sex-biased towards males.

### Expression of hippocampal IL-33 and its receptor peak following the second postnatal week, impacting ECM remodeling in the MyD88-cKO model

Thus far, we eliminated several mechanisms through which cKO microglia might impact inhibitory circuit development leading to increased GABAergic signaling in the adult hippocampus. For example, cKO microglia are not less efficient at inhibitory synaptic removal, and increased MyD88-independent signaling does not phenocopy brain and behavior outcomes. We next hypothesized that cKO microglia may be differentially interacting with ECM in the developing hippocampus. While the involvement of ECM in the development of primary sensory cortices has been previously well-characterized, recent work has described the role that microglia play in dentate gyrus ECM remodeling, with consequences for memory precision (Nguyen et al., 2020). Experience-dependent release of cytokine IL-33 from granule cell neurons in the dentate gyrus leads to increases in microglial interaction with aggrecan, a chondroitin sulfate proteoglycan (CSPG) in the ECM and a primary structural component of PNNs. This hippocampal-specific mechanism is particularly relevant because IL1RL1, the IL-33 receptor on microglia (also called ST2), signals through MyD88 (Figure 6A). Loss of IL-33 or its microglial receptor has been shown to impact excitatory/inhibitory balance and increase seizure susceptibility (Han et al., 2022). Therefore, we first determined whether the expression of cytokine IL-33 and its receptor IL1RL1 is developmentally regulated in the hippocampus, as has been shown in the thalamus (Vainchtein et al., 2018). There are two isoforms of IL-33 that are expressed in the adult hippocampus, *Il33a* from astrocytes and *Il33b* from neurons, allowing us to distinguish the source of IL-33 (Figure 6B) (Nguyen et al., 2020). We collected hippocampal tissue punches from wild-type mice from 5 ages across development (P7, P14, P20, P30, and P55, Figure 6C) and found that the expression of the neuron-derived *Il33b* isoform increased in early development and peaked in expression at P20 in males (Figure 6D). In contrast, mRNA expression of astrocyte-derived *Il33a* was not developmentally regulated in the male hippocampus (Figure 6D). We next quantified the expression of the receptor for IL-33, the IL1RL1 gene, *Il1rl1*, in the same tissue. This receptor is primarily expressed by microglia in the adult hippocampus (Nguyen et al., 2020). We found that *Il1rl1* was most highly expressed at P14 (Figure 6E). Interestingly, these expression patterns in the wild-type female hippocampus show less *Il33b*-specific upregulation in development (Figure S5A-B). Together, these findings provided a male developmental window of interest for probing microglia interactions with ECM, between P14-P20, when expression of both the ligand and receptor are at their highest.

**Figure 6.**
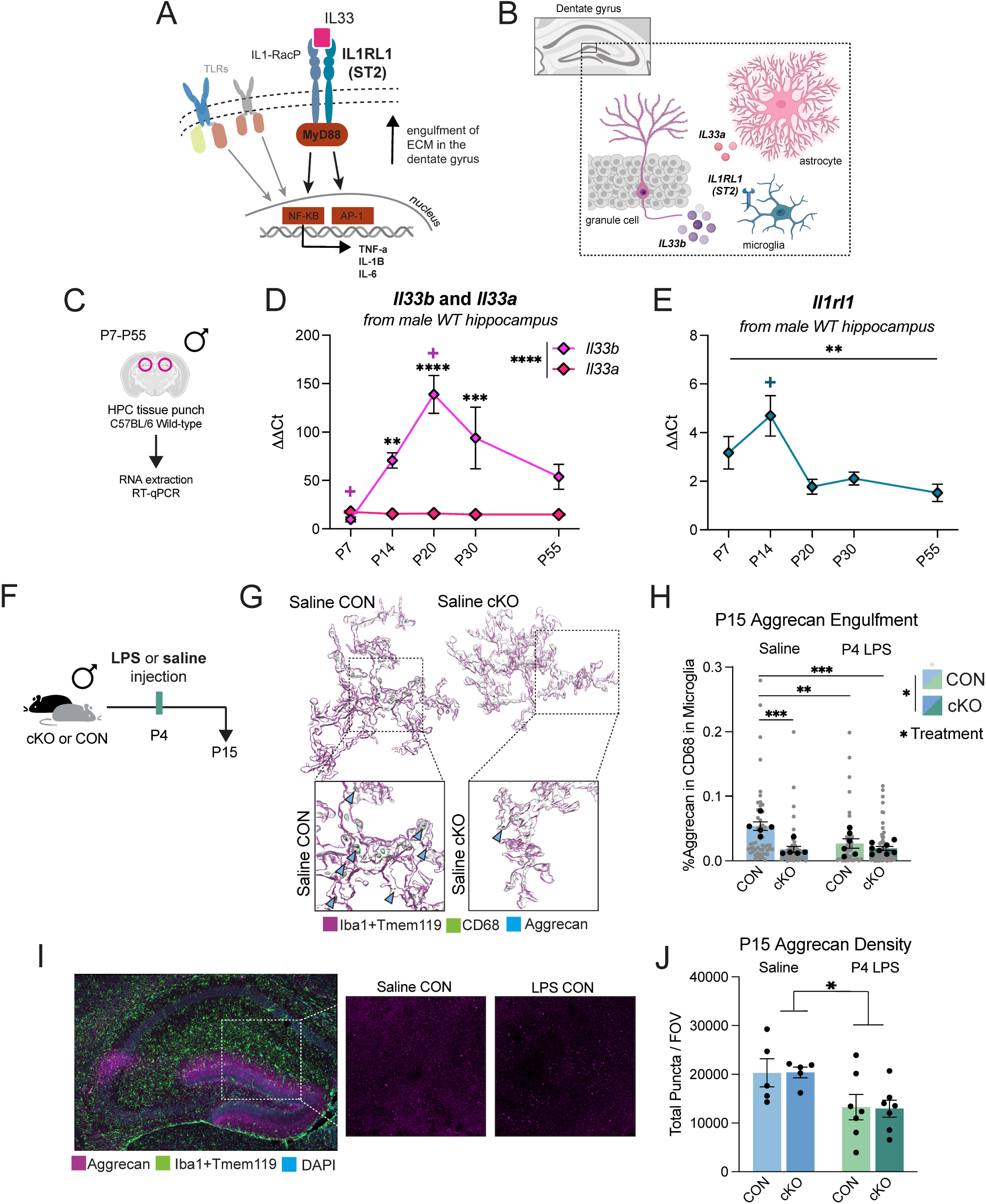
Expression of neuron-derived cytokine IL-33 and its receptor peak following the second postnatal week in the hippocampus, impacting ECM remodeling in the MyD88-cKO model **A.** MyD88-dependent signaling downstream of the IL-33 receptor, IL1RL1. **B.** Schematic of cellular sources of IL-33 isoforms, *Il33a* (astrocytes) and *Il33b* (neurons) in the dentate gyrus. **C.** Experimental design for RT-qPCR developmental timecourse in the male wild-type hippocampus. **D.** Developmental expression of *Il33a* and *Il33b*, significant main effects of age, isotype, and interaction by 2-way ANOVA. Asterisks indicate significant Bonferroni post-hoc between isotypes, + indicates significant Bonferroni post-hoc against adult expression (P55). **E.** Significant by one-way ANOVA, + indicates significant Bonferroni post-hoc against adult expression (P55), n = 8 mice/age. **F.** Experimental design for male microglial aggrecan engulfment after P4 endotoxin. **G.** Representative 3D reconstructions of microglia (magenta), lysosomes (green), and aggrecan (blue). **H.** Aggrecan engulfment at P15, significant main effects of treatment, genotype, and interaction effect by 2-way ANOVA. Asterisks indicate significant Bonferroni post-hoc, n = 5-7 mice/group. **I.** Representative image of aggrecan (magenta) in the dorsal hippocampus. **J.** Density of aggrecan in microglia images from the dentate gyrus (60x magnification), significant main effect of treatment by 2-way ANOVA, n = 5-7 mice/group. All error bars = SEM, all statistics run on individual animals (black or black-outlined circles), grey points represent individual microglia analyzed.

Because our previous experiments showed no changes in lysosomal content between genotypes by P18, we chose to quantify ECM remodeling in dentate gyrus microglia at P15, when cKO microglia contain more CD68 (Figure 2D-E). We utilized the same P4 LPS experimental paradigm (Figure 6F) and labeled microglia, lysosomes, and aggrecan in the dentate gyrus to volumetrically reconstruct microglia and their engulfment of ECM (Figure 6G). We found that P15 control microglia contained significantly more aggrecan in their lysosomes than any other group, and cKO microglia contained significantly less aggrecan compared to CONs across both treatment groups (Figure 6H). P4 LPS-exposed microglia, which are likely “primed” by early life inflammation, overall engulfed less aggrecan. However, when we quantified total aggrecan at this age, there was a main effect of P4 LPS treatment on the density of aggrecan (Figure 6I-J), suggesting that the decreased engulfment by LPS-treated microglia may be due to a global reduction in the aggrecan available to engulf, and, importantly, loss of MyD88 in microglia alone does not impact initial aggrecan deposition. Together, these findings support the hypothesis that normal developmental remodeling of aggrecan by microglia during postnatal hippocampal development is impacted by loss of MyD88-dependent signaling, likely through interruptions of IL-33-IL1RL1 ligand-receptor signaling.

### Bidirectional modulation of IL-33 early in life influences dentate gyrus microglia aggrecan remodeling in a MyD88-dependent manner

Having determined that cKO microglia engulf less aggrecan during developmental ECM remodeling (Figure 6H), we hypothesized this was due to an inability of cKO microglia to respond to IL-33. Microglial response to developmental IL-33 administration has been shown in other brain regions, such as the thalamus, to robustly induce classic macrophage activation genes and extracellular detection pathways (Han et al., 2022). To directly test if MyD88-cKO microglia can respond to IL-33 *in vivo*, we administered recombinant IL-33 (rIL-33) or sterile PBS control by intracerebroventricular (ICV) injection at P15 to CON and MyD88-cKO animals (Figure 7A). Four hours after injection, one hemisphere was taken for immunohistochemistry and the other for RT-qPCR of microdissected hippocampus. ICV injection allowed for diffusion of the cytokine to the hippocampal tissue without the confound of a direct injection injury and microglial reactivity at the site of interest. IL-33 activates IL1RL1 on microglia, which then signals through MyD88 to activate NF-KB-dependent cytokine release (Figure 6A). Administering IL-33 to CON mice induced a robust increase in *Tnf* and *Il1b*, which was blocked by loss of microglial-MyD88 in both male and female cKO mice (Figure 7B-C). When we looked at microglial morphology 4 hours after ICV injection, CON mice treated with IL-33 had a bushy, de-ramified morphology (Figure 7D-F). However, cKO microglia treated with IL-33 looked like PBS control-treated microglia and had significantly more Sholl intersections than IL-33-treated controls (Figure 7D-F). We next determined if developmental administration of IL-33 impacted microglial lysosomal content and aggrecan engulfment. We reconstructed microglia from IL-33 or PBS control treated MyD88-CON and cKO mice 4 hours after ICV injection and found that CON microglia responded to IL-33 by increasing their lysosomal volume and engulfing more aggrecan (Figure 7G-I). Again, loss of MyD88 in microglia prevented this increase in aggrecan phagocytosis (Figure 7G-I). These data confirm that MyD88-cKO microglia do not respond to IL-33 *in vivo*, and support previous literature that IL-33 increases aggrecan engulfment in adulthood <u>(Nguyen et al., 2020)</u>. While this experiment used likely supraphysiological levels of developmental IL-33, we know that the expression of neuron-derived IL-33 is highest between P14-20 in the homeostatic brain (Figure 3D). Finally, to functionally implicate a loss of developmental IL-33 signal to the changes we see in our cKO model, we virally delivered an shRNA against *Il33,* or a scrambled RNA (scrRNA) control, into the hippocampus of P4 wild-type mice (Figure 7J). At P15, microglia from scrRNA and shRNA P4 treated mice were reconstructed in the dorsal hippocampus. Viral expression was confirmed by GFP protein (Figure 7K). Reduction in *Il33* expression in early development led to a reduction in lysosomal volume and aggrecan engulfment by microglia at P15 (Figure 7L-M), in support of IL-33 signal in the dentate gyrus at P15 directing developmental aggrecan remodeling by microglia.

**Figure 7.**
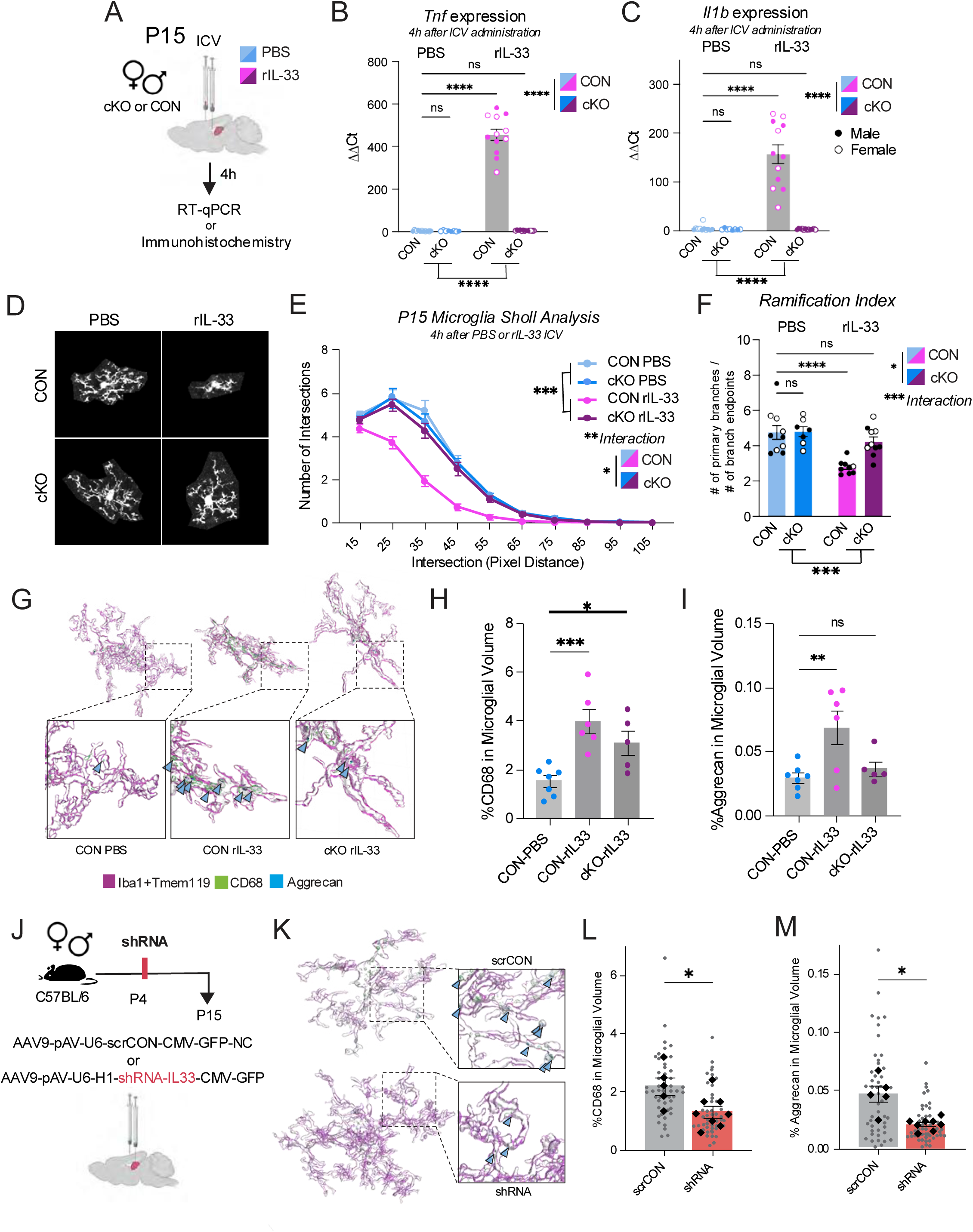
Acute IL-33 administration at P15 influences dentate gyrus microglial aggrecan remodeling in a MyD88-dependent manner **A.** Experimental design for intracerebroventricular (ICV) administration of recombinant (r)IL-33 or PBS control at P15 for RT-qPCR and IHC endpoints. **B.-C.** Expression of *Tnf* and *Il1b* 4 hours after ICV, significant main effect of treatment and interaction effect by 2-way ANOVA. Asterisks indicate significant Bonferroni post-hoc, n = 8-13 mice/group. **D.** Representative images of individual microglia after rIL-33. **E.** Sholl analysis of microglia, 3-way repeated measures ANOVA significant effects of treatment, genotype, intersection distance, and interaction effects (genotype x treatment). **F.** Ramification index (primary branches/endpoints), significant main effects by 2-way ANOVA of genotype, treatment, and interaction effect. Asterisks indicate significant Bonferroni post-hoc, n = 8-10 mice/group. **G.** Representative 3D reconstructions of microglia (magenta), lysosomes (green), and aggrecan (blue). **H.-I.** Lysosomal volume within microglia (**H.**) and aggrecan within lysosomes (**I.**), significant one-way ANOVAs, asterisks indicate significant Bonferroni post-hoc, n = 5-7 mice/group. **J.** Experimental design for short-hairpin (sh)RNA viral expression experiment. Mice were injected with scrambled (scr) control virus or shRNA against *Il33* into the hippocampus at postnatal day (P)4, and brains were collected at P15. **K.** Representative 3D reconstructed microglia from the dentate gyrus. **L.-M.** Lysosomal volume of scrCON and shRNA microglia (**L.**) and aggrecan engulfment within lysosomes within microglial volume (**M.**), significant by unpaired t-test. All error bars = SEM, all statistics run on individual animals (black or black-outlined symbols), grey points represent individual microglia analyzed.

## Discussion

Our study demonstrates that microglial MyD88 signaling serves as a critical regulator of inhibitory maturation in the developing hippocampus through control of cytokine responses, ECM remodeling, and synaptic refinement. Loss of MyD88 in male microglia reprograms microglial function, leading to altered interactions with inhibitory synapses, subsequent changes in their dynamics, long-term impacts on PVIs, and behavior deficits in a sex-specific manner. This work highlights a novel developmental role of IL-33 signaling through microglia in the dentate gyrus. The intersection of developmental alterations in innate immune signaling and PVI development/inhibitory maturation has been highly implicated in clinical observations, and here we propose one cellular mechanism through which these disease relevant changes may emerge.

This work began as an attempt to build a model of broadly blunted immune responsiveness in microglia through MyD88 deficiency. While loss of MyD88 led to the hypothesized decrease in cytokine response to inflammatory challenge, we were surprised to observe dramatic gene expression changes between CON and cKO microglia in the absence of any inflammatory challenge. This altered baseline gene expression is likely underlying later observed phenotypes, in particular the increased and more efficient phagocytosis observed in cKO males (Figure 2, Figure S2). Critically, the observed changes to male cKO gene expression patterns related to proteolysis that served as an initial hint to investigate ECM changes in this model fit with our later findings. We determined that MyD88 cKO microglia cannot respond to the IL-33 signal that induces developmental ECM engulfment, leading to over-deposition of aggrecan and PNNs in the dentate gyrus. One hypothesis to explain the downregulation by male cKO microglia of *inhibitors* of proteolysis and peptidases is that due to lack of remodeling signals from their environment due to MyD88 signaling loss, cKO microglia respond by attempting to increase the alternative, enzymatic breakdown of ECM. Future experiments could determine if specific downregulated gene targets (such as reduced expression of StefinA family genes *Stfa1*, *Stfa2l1*, *Stfa2*, and *Stefa3*, or other cystatin domain-containing genes like *Ctsdc4* and *Ctsdc5*, all of which inhibit cathepsins, thus increasing cathepsin activity) do indeed impact cKO microglia cathepsin activity *in vivo* or in slice, compared to CONs. This could determine if these gene expression changes specifically underlie the increase in phagocytic activity, and perhaps causal to the aberrant over-consumption of inhibitory synapses observed at P12. The other major observation from the sequencing results were the increase in type I interferon signaling in both male and female cKO microglia. We confirmed that administration of early life poly I:C to stimulate MyD88-independent signaling in the first postnatal week does not to reproduce the same behavioral and brain phenotypes observed in male MyD88-cKO mice (Figure S4). This, however, does not fully rule out involvement of MyD88-independent signaling in our findings, and further work is required to un-tease the impact of these signals, particularly in the early postnatal window.

The surprising finding that cKO microglia engulf specifically more inhibitory synaptic material at P12, but only cKO males end up with more PVI synapses later in life presents many interesting questions. First, why are PVI synapses spared? We confirmed that cKO microglia do not express a different abundance of GABA_B_ receptors (Figure S1). We believe the most likely answer is that the lack of ECM remodeling early in life leads to a physical barrier protecting microglia from reaching PVI synapses specifically. PVI synapses are known for their distinct peri-somatic distribution, and in the dentate gyrus (as in other brain regions), PVIs will synapse onto other PVIs. PVI somas are often surrounded by dense ECM in the form of PNNs, which we know are altered in cKO males (Figure 4). Perhaps as microglia attempt to degrade or remodel ECM by enzymatic release, which they are known to do in development (and is likely happening more in our model, as hypothesized by our sequencing findings), more inhibitory synapses are briefly exposed for remodeling at P12. Alternatively, as previously discussed, there appears to be a trending delay in the GABA switch in our cKO male mice (Figure S3), which could lead to a hyperexcitable state. PVIs are the most competent GABAergic cells to combat an over-excited system through their peri-somatic synapses, which exhibit great inhibitory control over their targets at the site of depolarization signal integration. The abundance of Syt2 synapses at P18 could simply be compensatory following when the GABA switch should have occurred but is instead delayed in cKO males. Alternatively, the decreased engulfment of aggrecan at P15 by cKO microglia could lead to a greater stabilization of later inhibitory synapses. The developmental delay in the GABA switch observed in cKO mice may also be influenced by the under-remodeling of the ECM by cKO microglia at P15; in the dentate gyrus, we would expect KCC2 levels to start rising during the second postnatal week, when ECM is being actively remodeled, impacting PVI maturation and activity, and therefore the GABA switch. However, whether the increased synaptic density or the increase in ECM is causal to the other is yet to be determined. These data highlight a sensitive window of development, when consequences of disruptions to homeostatic synapse and/or ECM remodeling are prolonged.

The discovery that PV interneuron density is increased in cKO adults is counter to much existing literature on the impact of early life challenges on PVI health, function, and long-term survival. However, there were a few avenues through which we hypothesized that microglial mechanisms may be involved in this counterintuitive increased neuronal survival. First, microglia are highly involved in the early postnatal cell death period, engulfing both apoptotic cells and living precursor cells, as well as releasing factors to induce apoptosis (Cunningham et al., 2013; Marín-Teva et al., 2004; Wakselman et al., 2008). Loss of MyD88 signaling in microglia did not impact cell density during the early postnatal interneuron apoptotic window (Wong et al., 2018) (Figure 4H-J), despite observed changes in cKO microglial phagocytosis. PV density is also influenced in the second week of life; work in the somatosensory cortex has implicated excitatory signaling from pyramidal neurons onto PVIs as highly influential to their survival, and that this influence is bidirectional (Wong et al., 2018). While circuit organization and development in the hippocampus are likely different, this literature does provide clues as to the importance of synaptic refinement in shaping PV cell survival and maturation, as a “better connected” PV cell is more likely to survive. While we do not see a change in VGlut2+ synapses at P18 in the cKO hippocampus, quantification of VGlut1+ synapses, or even synapses onto PV cells themselves, could help us better understand the connections between synaptic refinement and PV cell density.

However, we did find that there were increases specifically in Syt2+ synapses at P18. This is around the same time that PNNs begin to get more heavily deposited around PV cells in CA1 (Ramsaran et al., 2023). Super resolution imaging has also shown that at P30, the primary synapse type found within PNNs are Syt2+ synapses, which makes sense as most perisomatic synapses are from PV cells (Sigal et al., 2019). PNN deposition is also commonly used as an indicator of mature PV cells. Quantification of PV cell density by protein immunohistochemistry is confounded by the plasticity of the PV protein, which is known to be up-and down-regulated in response to cellular activity. Therefore the “increase” in PV density could simply be reflecting an increase in PV activity. A more active PV cell may have more Syt2+ connections (as seen at P18 and P30), and the GABAergic signaling onto surrounding cells may be stronger (as seen at P30 by electrophysiology). The increase in PNN+ PV cells seen in adulthood would also support this idea.

One interesting pattern that emerged across this work is the narrowing of broad effects to a specific vulnerable group over the course of maturation. Early in development, the loss of MyD88 from microglia impacted both males and females, and early life LPS exposure had little impact on these phenotypes. Next, from P18 to P30, only male cKO mice continued to exhibit more dramatic changes, such as in synapse abundance and electrophysiology. Finally, in adulthood, LPS treatment had the greatest effect on NOL behavior, more than half of the LPS cKO mice fell below the preference threshold, while only half did in the LPS CON group. In the case of social preference, the major behaviorally deficient group was the male LPS cKOs. While it is difficult to implicate a specific cellular mechanism in this behavioral change, PV cells are known to be critically important for discrimination functions (van Dijk and Fenton, 2018). One theory, then, might be that more PV cells lead to better performance on this task. This, however, was not the case, leading us to investigate another known contributor to memory precision in the hippocampus: the ECM. It was recently shown that proper PNN maturation in the hippocampus is critical for memory behaviors, and the key window for this to occur is the very beginning of the fourth postnatal week, the same time when our deficits begin to become P4 LPS cKO specific (Ramsaran et al., 2023). While PV cell function was shown to be necessary for memory precision, modulating PNN deposition and maturation was sufficient to induce behavioral deficits, indirectly through impacts on PV cells, which we suspect is more likely the cellular contributor to these behavior changes. Interestingly, previous work characterizing IL-33 knockout mice similarly identified deficits in social discrimination, but not sociability (Dohi et al., 2017).

One limitation of this work is the use of a constitutive MyD88-cKO model. Although an inducible transgenic approach to remove MyD88 from microglia would in theory allow for more precise temporal conclusions, efficient recombination in microglia during the first postnatal week, when MyD88-IL-33 signaling is critical in the hippocampus, remains technically impossible with the currently available tools (Faust et al., 2023). While we cannot entirely rule out contributions of microglial MyD88 in embryonic interneuron development, our data suggests that at least PV and STT interneurons appear in normal density prior to P4.

Together, our findings provide a mechanistic link between an immune cue, cytokine IL-33, and the closure of a developmental plasticity window in the hippocampus via microglia-ECM interactions, rather than simply through direct synaptic interactions. The inability of MyD88-cKO microglia to respond to IL-33 prevented proper developmental ECM remodeling, leading to aberrant PNN stabilization and long-lived changes in inhibitory system function. This work identifies a novel developmental application of an immune signaling pathway that provides insight into how early life inflammatory events may bias neurodevelopmental trajectories in a sex-specific manner.

## Methods and Materials

### Animals

All experiments were conducted in accordance with the NIH *Guide to the Care and Use of Laboratory Animals* and approved by the Duke University Institutional Animal Care and Use Committee (IACUC). Animals were group housed in a standard 12-12h light/dark cycle. MyD88-floxed mice were obtained from Jackson Laboratory (Stock #008888) and crossed to Cx3cr1-BT-Cre (MW126GSat), which were provided by L. Kus (GENSAT BAC Transgenic Project) until all offspring were fully MyD88 floxed (F/F) and either Cre-negative (Cre^0/0^ or CON) or Cre-positive (Cre^tg/0^ or cKO). Mice were then backcrossed onto a fully Jackson background. The *Cre* transgene was maintained in males for all experimental studies whenever possible. Each litter was bred to contain a mixture cKO offspring and CON offspring to be used as littermate controls. CON and cKO experimental mice were co-housed for all experiments. For all experimental endpoints, mice from at least 3 different litters were included in each sex/genotype/treatment group, unless otherwise noted.

### Genotyping

Genotyping of transgenic animals was conducted from tail (at postnatal day (P)14) or toe (at P4) DNA using polymerase chain reaction (PCR). For all experimental animals, genotyping confirmed fully floxed MyD88 alleles as well as the absence or presence of Cre recombinase (<u>Table S1</u>). High recombination efficiency has been previously validated in this mouse line by PCR followed by gel electrophoresis (Rawls et al., 2025).

### Microglia isolation

Male and female CON and cKO mice were sacrificed by intraperitoneal (i.p.) injection of Avertin (tribromoethanol, if age <P14) or CO_2_ inhalation and then transcardially perfused with ice-cold saline to eliminate blood immune cells in brain vasculature. Microglia were isolated from brain tissue using CD11b magnetic beads as previously published (Bordt et al., 2020). Briefly, bilateral hippocampi were dissected out on ice then enzymatically digested (1.5 mg/mL collagenase A, 0.4mg/mL DNAse I in Hank’s Buffered Salt Solution (HBSS)) in 15-minute increments in a 37°C water bath between stepwise mechanical dissociation through pulled glass Pasteur pipettes. Dissociated samples were run through nylon filters and spun to pellet cells at 1,200 rpm at 4°C for 10 minutes. Cells were then incubated with Miltenyi Biotec MACS CD11b antibody-conjugated magnetic beads for 15 minutes on ice, washed, then passed through magnetic beds columns to isolate CD11b negative (non-microglia) from CD11b positive (microglia) populations. After pelleting again, cells were washed and then re-suspended in appropriate solution for desired endpoints.

### Ex-vivo cultured microglia stimulation and MSD supernatant quantification

After CD11b bead isolation of microglia from P4-6 male and female CON and cKO hippocampus and cortex, CD11b+ cells were plated at a density of 50,000 cells/well in a v-bottomed 96-well plate in microglia culture media (100 u/mL penicillin-streptomycin, 1mM sodium pyruvate, 2mM L-glutamine, 4.2μm/mL Forskolin, and 1X N2 Media Supplement in DMEM). Wells were treated with either 100ng/mL lipopolysaccharide (LPS) or equal volume of 1X PBS for 4 hours in a 5% CO_2_, 37°C cell culture incubator. Cells were then spun at 500 g for 2 minutes, and then supernatant was removed and stored at-80°C until time of assay. A total of 3-6 wells/animal/treatment were combined for supernatant analysis, and combined well samples from each animal were run in duplicate. Mesoscale Discovery (MSD) custom UPLEX panel of a custom set of biomarkers were performed on blinded samples by the Biomarkers Core Facility at the Duke Molecular Physiology Institute.

### RNA extraction for sequencing

Male and female CON and cKO mice were injected i.p. with saline or 330μg/kg LPS at P14, then returned to their home cage for 2 hours. Mice were then transcardially perfused with ice-cold saline and hippocampi were bilaterally dissected. Microglia were isolated as described above. At the final step, pelleted CD11b+ microglia (and/or CD11b-, non-microglia) were resuspended in 500μL Trizol and stored at-80°C until RNA extraction. Sequencing samples were all processed across 2 days together in small batches encompassing all experimental groups.

RNA was extracted from pellets homogenized in Trizol by addition of chloroform (1:5 to Trizol volume), vortexing for 2 minutes, then centrifuging at 11,800 rpm for 15 minutes at 4°C. The clear aqueous phase was moved to a new tube, and then RNA was precipitated by vortexing then centrifuging with Glycogen (1μL) and isopropanol (∼1:1 with aqueous phase). Supernatant was then removed, and pellets were washed and spun twice with ice cold 75% ethanol, then dried and resuspended in RNAse-free water.

### Bulk RNA-sequencing and analysis

Sample quality was verified, then libraries were constructed for sequencing on an Illumina NovaSeq 6000. RNA quality was assessed by the Duke University School of Medicine Sequencing and Genomic Technologies (SGT) core facility, and the best RIN value samples were selected for library preparation. Sequencing libraries were constructed by SGT using the Illumina Stranded mRNA-Seq HyperPrep Kit with 50 ng total RNA input per sample. Libraries were pooled and sequenced on an Illumina NovaSeq 6000 to generate 2 x 100 bp paired-end reads, yielding ∼28–120 million paired-end reads per sample with ≥ 93% of bases ≥ Q30. Samples with a PF yield below 14 Gb were excluded from further analysis: two samples were automatically removed due to PF yields below 10 Gb, and two additional samples were excluded at a later stage because they expressed very low levels of microglia-specific genes (Cx3cr1, Tmem119, P2ry12). Raw sequencing reads were assessed for quality using FastQC and adapter sequences and low-quality bases were trimmed using Trimmomatic. Cleaned reads were aligned to the Mus Musculus GRCm38 mm9 genome using STAR. Read counts were generated using featureCounts, non-protein-coding genes were filtered out, and only genes that had 1 count or more in at least 4 samples were used for differential gene expression analysis using DESeq2 (Love et al., 2014). Samples were first filtered by sex and treatment, then differentially expressed genes between genotypes were identified using an adjusted p-value cutoff of 0.1. Volcano plots of up-and down-regulated genes were generated using a custom script, which can be found at github.com/dzjulia. DAVID bioinformatics resource was used to generate Gene Ontology figures (Huang et al., 2009; Sherman et al., 2022).

### Immunohistochemistry

Mice <P14 were sacrificed by intraperitoneal (i.p.) injection of Avertin, and mice P14+ were sacrificed by CO_2_ inhalation. Mice were then transcardially perfused with ice-cold saline followed by 4% paraformaldehyde (PFA) and brains post-fixed in 4% PFA for 24 hours at 4°C. Brains were then dehydrated in a 30% sucrose in 1X PBS solution with 0.1% sodium azide for at least 72 hours at 4°C. Brains were flash-frozen in 2-methylbutane pre-chilled on dry-ice and stored at - 80°C until cryosectioning at 40μm into tubes of cryoprotectant. Tissue was stored at-20°C until use.

For all endpoints excluding synapse density, free floating tissue sections were wash 3 times in 1X PBS at room temperature (RT) on a shaker for 10 minutes, then blocked in 5% or 10% normal goat serum (NGS) and 0.1 or 0.3% Triton-X in 1X PBS for 1-2 hours at RT on a shaker. Tissue was then moved into primary solution, which matched blocking solution with the addition of primary antibodies (see <u>Table S2</u>), for an overnight incubation at 4°C. Primary solution was then washed off by washing 3 times in 1X PBS at RT on the shaker for 10 minutes. Tissue was incubated, covered, on a shaker at RT for 2-4 hours with secondary antibodies in 1X PBS (see <u>Table S2</u>), then washed 3 times again before mounting on subbed slides and coverslipping with Vectashield Plus with DAPI or Fluoromount G with DAPI.

### Microglia reconstruction imaging and analysis

Images were acquired on an Olympus FV3000 inverted confocal. Individual cells were imaged using a 60X magnification with 1.3x optical zoom, capturing the full depth of tissue using a 0.33μm step Z-stack. Images were converted to IMARIS files (v9.5), and “Surfaces” tool was used to render the volume of individual cells, and surface masking was used to reconstruct CD68 within cell volume. Phagocytic capacity was calculated as (total volume CD68 within microglia volume / total volume of that microglia)*100. To quantify synaptic and aggrecan engulfment, surface masking of CD68 volume was used to determine the surface volume of synaptic or aggrecan material within CD68, and engulfment of material was calculated as (total volume of material within CD68 within an individual microglia / total volume of that microglia)*100. All reconstructions were performed blinded to experimental condition. At least 4 images were captured per animal, and at least 6 cells/animal were included in all analyses. All statistics were run on biological replicates, which were comprised of the average of all cells for that animal. An equal number of microglia from within the hilus and in the molecular layer of the dentate gyrus were included in all analyses.

### Neonatal LPS injections

To administer LPS or saline at postnatal day 4, pups were injected one at a time to minimize time away from the dam. Pups were briefly removed from the nest and weighed, then placed on a clean towel, where they were injected subcutaneously with 330μg/kg LPS dissolved in 0.9% sterile saline, or 0.9% sterile saline, with a 30-gauge needle to minimize leaking. Tails were then marked with a sharpie before returning to the nest to ensure the same pup was not injected more than once. The total duration of maternal separation for each pup was between 30 seconds to 1 minute. All mice within one litter received the same treatment to prevent indirect exposure to the alternate drug treatment and to avoid differential maternal care. The timing and dose of injection was selected based on previous work from the lab (Bilbo et al., 2005b, 2005a; Bland et al., 2010; Schwarz and Bilbo, 2011; Smith et al., 2020; Williamson et al., 2011)

### Microglia 2-D sholl analysis

Images were acquired on an Olympus FV3000 inverted confocal. Tilescans of the dorsal hippocampus were captured using 20X magnification, and 5 z-stacks were imaged at 1μm step size. The microglia channel was z-projected and individual cells from the dentate gyrus (at least 6 per image, at least 2 images/animal) were selected by an experimenter blinded to conditions. Those individual cells were then automatically pixel segmented using the Ilastik software (Berg et al., 2019) after training on a random subset of 6 cells. The output binary segmentations of each cell were then converted to 8-bit and then run through a custom python script to skeletonize the images and perform sholl analysis (calculate intersections of skeletonized cell processes on concentric circles surrounding the cell, 10 pixel distance apart from one another). Ramification index was calculated as the ratio of primary branches (intersections at first ring) to branch endpoints. Code for 2D sholl analysis can be found here: https://github.com/bendevlin18/sholl-analysis-python

### Microglia density quantification

Images were acquired as tilescans of the dorsal hippocampus captured on a Zeiss Axio Imager Z1 epifluorescent microscope with apotome attachment using a 10X objective. Maximum intensity projections were produced from 19-step Z stacks captured with a 1.67μm step size. Projected images were then manually cropped using FIJI to include only the dorsal hippocampus and are the total area measured. Cropped hippocampi images were then imported into Ilastik software, where pixel segmentation followed by object classification were used to identify and count individual Iba1+ somas, which were then normalized to total area of each dorsal hippocampus.

### RNA-FISH *in-situ* hybridization for GABA_B_ receptor subunits

RNA-FISH was performed using Molecular Instruments Multiplexed HCR Immunofluorescence (IF) + RNA-FISH protocol for free-floating fixed tissue sections. P12 hippocampal sections were collected and prepared as described for immunohistochemistry, blocked in 5% normal donkey serum and 0.1% Triton-X in PBS, and then protein and RNA detection protocols were performed following the manufacturer’s instructions. Chicken anti-Iba1 primary antibody (Synaptic Systems, Cat# 234009, 1:500) was used for IF with Donkey anti-chicken HCR probe, and custom probes were designed for *Gabbr1* and *Gabbr2* (Table S4). Images were acquired on an Olympus FV3000 inverted confocal. Individual cells were imaged using a 60X magnification with 1.3x optical zoom, capturing the full depth of tissue using a 0.33μm step Z-stack. Images were converted to IMARIS files (v10.0), and “Surfaces” tool was used to render the volume of individual microglia. “Spots” tool was then used to standardized size (0.4μm) and quality (170) settings, and number of *Gabbr1* and *Gabbr2* puncta were calculated. Percent of individual microglia with both puncta were calculated, as well as the ratio of *Gabbr1* to *Gabbr2* puncta per cell. Sections from at least 3 animal/group were pooled for analyses, so data for only this experiment was analyzed on a per cell basis, without individual biological replicate information.

### qPCR

Wild-type (C57BL/6J) mice were sacrificed and saline-perfused at 5 ages between P7-P55 and brains were flash-frozen then stored at-80°C. Hippocampus was punched from frozen brains in a cryostat then directly placed in Trizol for RNA extraction (as described above). RNA concentration in each sample was determined by Nanodrop spectrophotometer and 1000ng of RNA was used to create cDNA by Qiagen QuantiTect Reverse Transcription kit. RT-qPCR was run using a QuantStudio Real-Time PCR system by in-house primers (Table S1) and Sybr. All samples were loaded in duplicate and run on the same day by the same experimenter to reduce variability. 18S served as the control sequence for sample normalization and fold change was calculated using the 2^−ΔΔCT^ method.

### Aggrecan quantification

The same images captured for aggrecan engulfment endpoints were used to quantify aggrecan puncta within the same field of view. The “Spots” function on IMARIS was used to count the aggrecan within each image, and the same settings were used for each image with auto thresholding. Values for each image were averaged so that each animal was represented by one value. The same pattern was observed using FIJI mean fluorescence intensity (data not shown).

### Synapse labeling

For all synapse density endpoints free floating tissue sections were washed 3 times in 1X TRIS-buffered saline with 0.2% Triton X-100 (TBS-Tx) at RT on a shaker for 10 minutes. Excitatory synaptic proteins were blocked in 5% NGS in TBS-Tx for 1 hour at RT on a shaker. Inhibitory synaptic proteins were blocked in 10% NGS in TBS-Tx for 1 hour at RT on a shaker. Tissue was then moved into primary solution, which matched blocking solution with the addition of primary antibodies (see <u>Table S2</u>). Excitatory proteins were incubated overnight at 4°C, while inhibitory proteins were incubated at 4°C for 48 hours. Primary solution was then washed off by washing 3 times in TBS-Tx at RT on the shaker for 10 minutes. Tissue was incubated, covered, on a shaker at RT for 2 hours with secondary antibodies in TBS-Tx (see Table S2), then washed 3 times again before mounting on gelatin-subbed slides and coverslipping with Fluoromount G with DAPI.

### Synapse quantification

All synaptic endpoints were imaged within 48 hours of mounting tissue whenever possible. Images were captured using an Olympus FV3000 inverted confocal microscope using a 60X magnification with 1.3X optical zoom. For each brain section, 2 x 2 tilescans were captured for each hemisphere, with the field of view centered over the hilus of the dentate gyrus, with 5 slices captured at a Z step size of 0.33μM, centered on the middle of the tissue thickness. Synaptic density was quantified using the FIJI Syn-Bot plug-in manual thresholding option (Savage et al., 2024).

### Electrophysiology

Male cKO and CON mice were deeply anesthetized using isoflurane before rapid decapitation and brain removal. 250 μm coronal sections were collected in ice-cold modified artificial cerebral spinal fluid (aCSF: 95% oxygen, 5% carbon dioxide, 194 mM sucrose, 30 mM NaCl, 4.5 mM KCl, 1 mM MgCl2, 26 mM NaHCO3, 1.2 mM NaH2PO4, and 10 mM D-glucose) using a Leica VT1200 vibrating microtome. Brain sections were then transferred to regular aCSF (95% oxygen, 5% carbon dioxide, 124 mM NaCl, 4.5 mM KCl, 2 mM CaCl2, 1 mM MgCl2, 26 mM NaHCO3, 1.2 mM NaH2PO4, and 10 mM D-glucose) and incubated at 32°C for 30 min for recovery. Slices were then removed from the incubator and stored at room temperature until recording.

Slices were hemisected, placed into a recording chamber, and perfused with temperature-controlled aCSF (29–31 °C) containing 50 μM DL-AP5 and 5 μM NBQX. Granule cells were voltage clamped at −60mV using a MultiClamp 700B Amplifier, and spontaneous inhibitory postsynaptic current (sIPSC) events were recorded using a borosilicate glass pipette (3–5 MΩ resistance) filled with chloride-based internal solution (150 mM CsCl, 10 mM HEPES, 2 mM MgCl2, 0.3 mM Na-GTP, 5 mM QX-314, 3 mM Mg-ATP, 0.2 mM BAPTA, and 2-4% biocytin).

Signals were filtered at 2 kHz, digitized at 10 kHz and acquired using Clampex 10.4.1.4 software from Molecular Devices, and data later analyzed using Mini Analysis Program (Synaptosoft).

### Immunoblotting

MyD88-cKO and CON mice were sacrificed between P28-P31, saline perfused, and bilateral hippocampi microdissected, flash frozen, and stored at-80°C until processing. Tissue was then homogenized using in N-PER extraction reagent with protease and phosphatase inhibitors and protein concentration normalized across samples. For NKCC1, KCC2, Syt2, and beta-actin, ∼50μg of protein of each sample was loaded into 4-15% Mini-PROTEAN TGX Stain-Free precast gel (Bio-Rad) and run in Tris/glycine/SDS running buffer (Bio-Rad). Proteins were transferred using Trans-Blot Turbo semi-dry transfer system to a PVDF membrane using the standard transfer setting. Membranes were then blocked with 5% milk in TBS-Tween20 for 1 hour before incubating overnight with primary antibodies (Table S3) diluted in blocking solution. The next day, primary was removed with TBS-Tween20 washes then incubated in secondary antibodies (Table S3) in 5% milk in TBS-Tween20 for 1 hour, covered. Images of membranes were captured on a ChemiDoc. Protein abundance was normalized to β-actin and quantified in FIJI using the Analyze > Gels tool.

### Adult interneuron and extracellular matrix quantification

Immunohistochemistry was performed as described above with appropriate antibodies (Table S2). Tilescans of the dorsal hippocampus were captured on a Zeiss Axio Imager Z1 epifluorescent microscope with apotome attachment using a 10X objective. Maximum intensity projections were produced from 19-step Z stacks captured with a 1.67μm step size. Projected images were imported into QuPath for manual blinded counting of PV cells and PNNs in each channel in isolated (Bankhead et al., 2017). Colabeled PV/PNNs were then quantified by PV/PNN label overlap. Cell number was normalized to the area measured in QuPath.

For WFA intensity quantification, dorsal hippocampus images used to count PV cells and PNNs were re-analyzed for this experiment. Maximum intensity projections of tilescans were produced and PV channel was used to identify the outlined area of the dorsal hippocampus. ImageJ measurement was used to measure the integrated density of WFA fluorescence within the dorsal hippocampal area for each image

### Neonatal interneuron quantification

Brains for RNAscope endpoints were collected from P0, P4, and P8 pups were following deep anesthesia from avertin injection and saline transcardial perfusion. Non-fixed brains were embedded in O.C.T. over dry ice and stored at-80°C until sectioning. Brains were then sectioned at 20μm and immediately slide-mounted on Superfrost Plus slides, and again stored at-80°C until RNAscope procedure. Manufacturer’s protocol (ACD Bio) was followed, consisting of a 15 minute 4% PFA fixation, 3 x 5 minute EtOH washes (50%, 70%, and 100%), and a 30 minute protease treatment. Tissue was then treated with the following primary probes, covered in Parafilm to avoid evaporation, for 2 hours in a 37°C oven: C2-*Lhx6*, C3-*Sst*. Tissue was then amplified and coverslipped with Vectashield plus DAPI. Images were captured as previously described for adult quantification, and QuPath was again used for blinded quantification as described, using the DAPI channel to distinguish individual cells.

### Sociability and Social Novelty Preference behavior

Social behavior was tested in male and female mice between P35-37. Mice were first habituated to testing room and the testing apparatus-1 day before testing. A separate 3-chambered arena was used for each sex to avoid distracting odor cross-contamination. All stimulus animals were WT strangers and were age-and sex-matched to the experimental animal. Smaller enclosures were centered in opposite ended chambers of the assay. For sociability testing, a stranger conspecific stimulus animal was placed in one chamber and an object (rubber duck) was placed in the other. Experimental mice were placed in the center chamber and their behavior was recorded for 5 minutes by an overhead camera in EthoVision XT software (Noldus) for trial management. At the conclusion on 5 minutes, the object was replaced with a new, novel stranger conspecific (now the “novel” animal), and the time spent interacting with the novel and “familiar” animal (from the original sociability trial) was recorded for 5 minutes. The arena was cleaned thoroughly between each experimental animal and the social vs. object side was alternated each trial. All videos were manually scored by a blinded experimenter to measure total time spent actively investigating either chamber. Sociability was scored as (time spent interacting with social stimulus / (time spent interacting with object + time spend interacting with social stimulus)) * 100. Social Novelty Preference was calculated as (time spent interacting with novel social stimulus / (time spent interacting with familiar social stimulus + time spend interacting with novel social stimulus)) * 100.

### Open Field behavior

Open field behavior was run ∼P50 to assess “anxiety-like” behaviors, as well as location measures. Mice were allowed to freely explore an open arena testing chamber (45cm x 45cm, opaque white walls) for 10 minutes after being acclimated to the testing room for 1 hour prior. Behavior was recorded from above, and Ethovision XT software (Noldus) was used to quantify distance traveled, average velocity, as well as time spent in the center (15 cm x 15 cm square) of the arena.

### Y-maze spontaneous alternation behavior

To test working memory, mice were allowed to freely explore the 3 arms of a y-maze (35 cm arm length, 5cm arm width, with 10 cm wall height, covered by plexiglass lids) for 5 mins. Behavior was recorded using an overheard camera and Ethovision XT software for quantification. Alternation percentage was calculated as [consecutive A, B, C arm visits, or “spontaneous alternation” / (total number of arm entries − 2)] x 100.

### Novel Object Location behavior

Novel object location (NOL) behavior paradigm was performed about P50. Mice were previously exposed to the testing chamber in the morning of testing day 1 for open field behavior. Approximately 2 hours after open field testing, mice were returned to the testing chamber for Trial 1. A square, solid color visual stimulus was added to 1 wall of the all-white, high-walled testing chamber to provide interior proximate visual localization cues. Two identical, uniquely shaped objects were secured to the base of the chamber, equally spaced and in the top 1/3 of the chamber. Objects were previously tested in non-experimental animals to ensure that mice were not fearful/avoidant of the objects. For Trial 1, mice were allowed to explore the familiar chamber and both objects for 10 minutes and were recorded with an overhead camera into Ethovision XT software (Noldus) for trial organization. Mice were then returned to their home cage and the colony room overnight, and Trial 2 was recorded the next morning, 20 hours after Trial 1. Mice acclimated to the testing room for 30 minutes prior to Trial 2, where one object was moved to the bottom 1/3 of the testing chamber (novel location) and the other was left in its original location (familiar location). Mice were again recorded for 10 minutes, and their interactions manually scored by a blinded experimenter. Time spent interacting with each object was documented in each trial, and trial scoring stopped when the mouse reached a criterion of 30 total seconds spent interacting with either object. Preference for the novel object location was calculated as (total time spent interacting with the novel location object / (time spent interacting with the novel location object + familiar location object)) * 100. Chambers and objects were thoroughly cleaned between each trial to avoid distracting odorants. When males and females were run on the same day, males were always ran first to avoid female scent distraction.

### Poly I:C administration

Poly I:C (Polyinosinic–polycytidylic acid potassium salt, Sigma-Aldrich, P9582) was injected subcutaneously from P2-P6 at a 5mg/kg dose dissolved in 0.9% sterile saline, or 0.9% sterile saline control, as previously described (Schwabenland et al., 2023). The same protocol was followed as described in “Neonatal LPS injections,” where pup separation from the dam is minimized as much as possible, and entire litters receive a single treatment in order to minimize treatment-induced changes in maternal care.

### Serum IL-6 Quantification

Following the final P6 poly I:C or saline injection, pups were sacrificed 4h later by i.p. injection of Avertin (tribromoethanol), and loss of response was confirmed by toe-pinch. Trunk blood was then collected by decapitation and serum prepared and then processed according to manufacturer’s instructions for the Mouse IL-6 ELISA (R&D Systems, M6000B).

### P15 IL-33 administration

CON and cKO adult mice were anesthetized using 3-4% isoflurane for the duration of surgery and breathing was closely monitored. Head fur was shaved, then mice were placed in a stereotax (Stoeling) and skin was cleaned with betadine and ethanol before incision. Coordinates were selected to reach each ventricle anterior to the hippocampal formation:-0.3 AP, +/-1.5 ML,-2.5 DV to bregma. Holes were drilled bilaterally into the skull and 1µL of sterile 1 X PBS or recombinant IL-33 (rIL-33, R&D Systems, 100μg/mL) reconstituted in sterile 1 X PBS was administered by Hamilton syringe to each ventricle sequentially at a rate of 0.25 µL per minute.

Mice received 5mg/kg ketoprofen prior to being removed from anesthesia. Brains were collected 4h following injection as previously described for RNA extraction, then one hemisphere was drop-fixed in 4% PFA and processed as previously described for IHC.

### P4 shRNA administration

Sequences for shRNA targeting of the *Il33* gene were selected based on previous work (Hatzioannou et al., 2020) Viral constructs were created and packaged by WZ Biosciences as follows: AAV9-pAV-U6-scrambledControl-CMV-GFP-NC and AAV9-pAV-U6-H1-shRNA against IL33-CMV-GFP. Virus was aliquoted and stored at-80°C, then kept on ice until injection. P4 intrahippocampal viral infection was performed as previously described at P1 (Devlin et al., 2024). In brief, mice were anesthetized in a conical tube placed in an ice bath for 10 minutes, then placed on an ice pack following toe-pinch confirmation of anesthesia. Using a glass pulled needle, mice were bilaterally injected with 100nL per hemisphere of virus at a rate of 250nL/min using a microinjector. Mice were placed on a warming pad until awake, then returned to their homecage. Brains were collected for immunohistochemistry as previously described at P15, sectioned in a cryostat at 40μm, then GFP signal was amplified by IHC (Table S2) and imaged across the brain to determine which animals to include in microglia analysis endpoints in the hippocampus. Sections from brains with validated hits by hippocampal GFP expression were then photobleached overnight to quench GFP signal, then IHC was performed and images captured and analyzed for microglial engulfment of aggrecan and aggrecan density as previously described.

## Acknowledgements

This work was supported by funding from the NIH – F31AA030712 to JED and 1U01-AA029969 to SDB – and the Charles Lafitte Foundation Program for Research in Psychology & Neuroscience at Duke University (to JED). Thank you to Janet Huebner from the Biomarkers Core Facility at the Duke Molecular Physiology Institute for assistance with the MSD panel and the Duke Light Microscopy Core facility for microscopy and analysis support. Thank you to Dr. Hannah Staley, Dr. Sushanth Kumar, and Dr. Juan Ramirez for training and guidance on microglia culture experiments. Special thanks to Dr. Marcy Kingsbury for conducting experiments that served as foundational preliminary data for this work, and to the Integrative Neuroscience Initiative on Alcoholism – Neuroimmune for their valuable insights on this project. The authors have no conflicting financial interests.

---

**Table S1:**
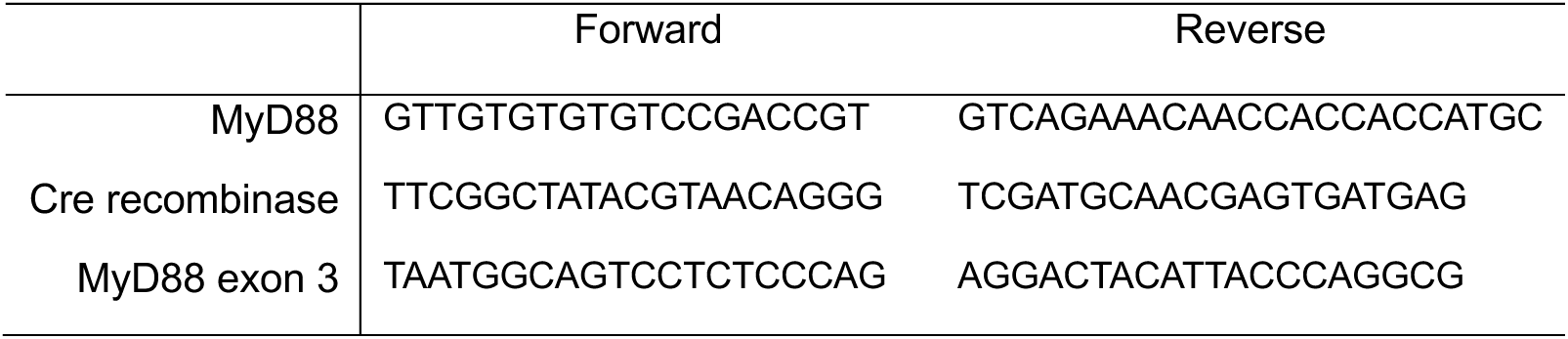
Genotyping Primer Sequences.

**Table S2:**
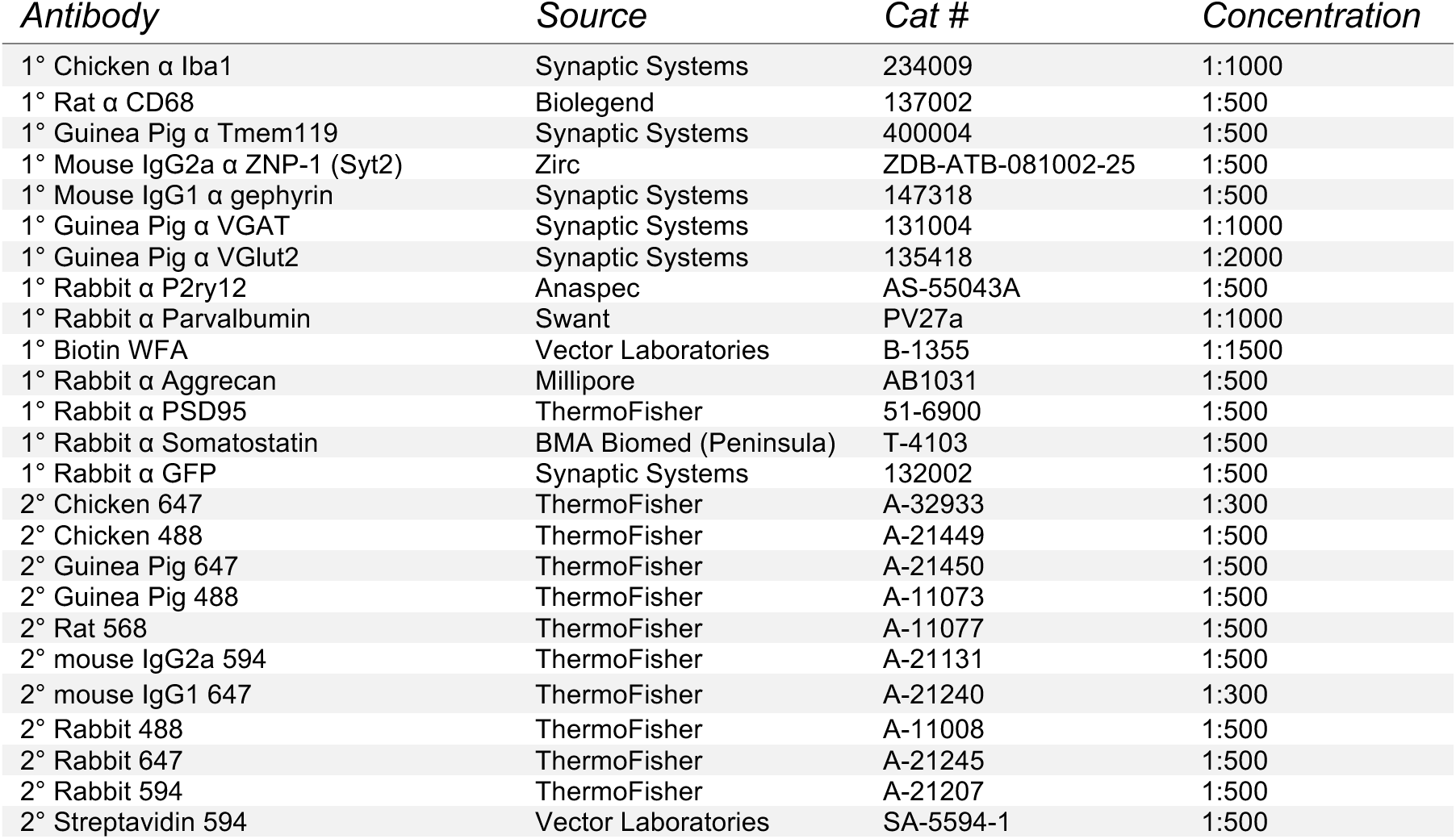
Immunohistochemical Antibodies.

**Table S3:**
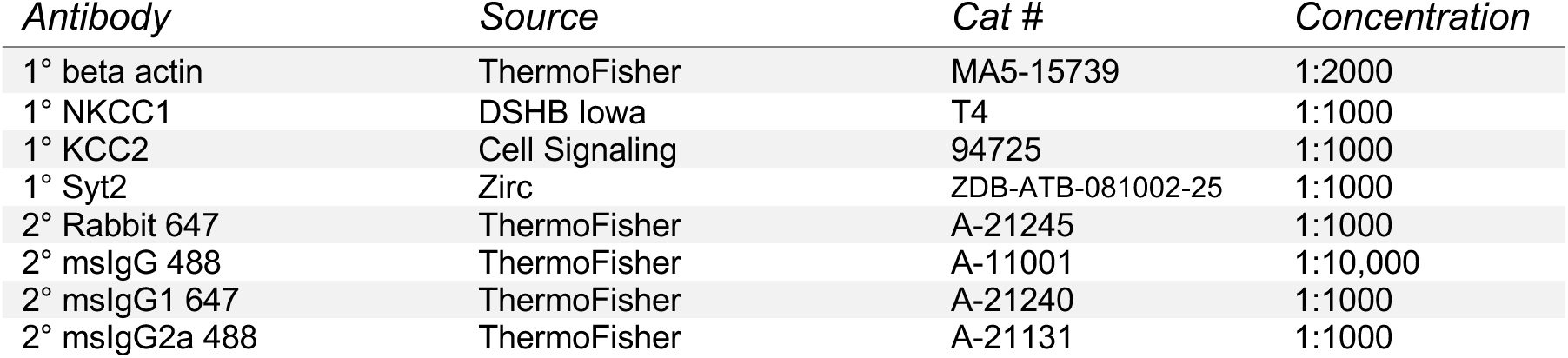
Immunoblot Antibodies.

**Table S4:**
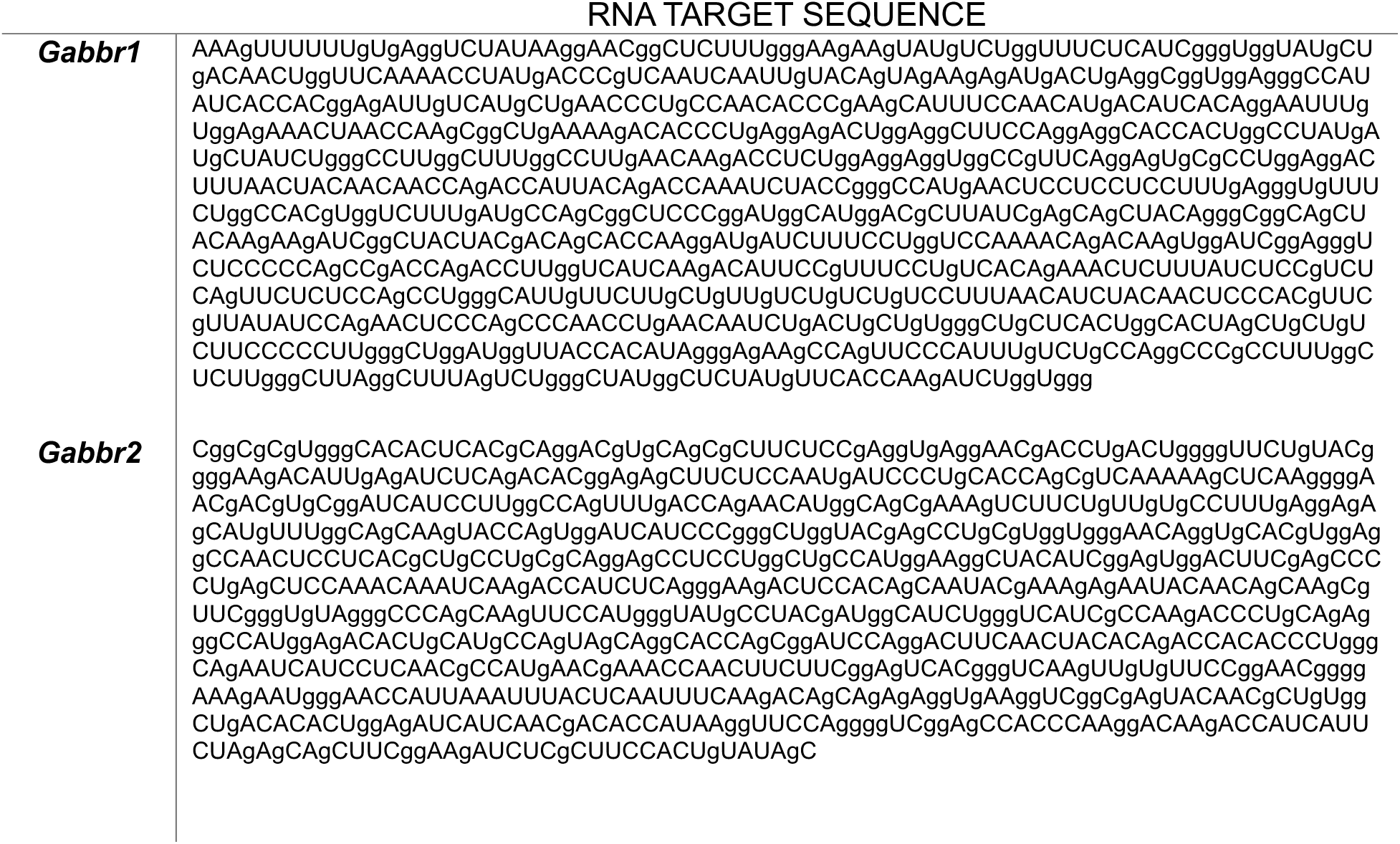
Custom Probes for RNA-FISH.

