## Supplemental Figures 1-5 for "Microglial MyD88-dependent signaling influences extracellular matrix development and interneuron maturation in the hippocampus"

Supplementary Figure 1

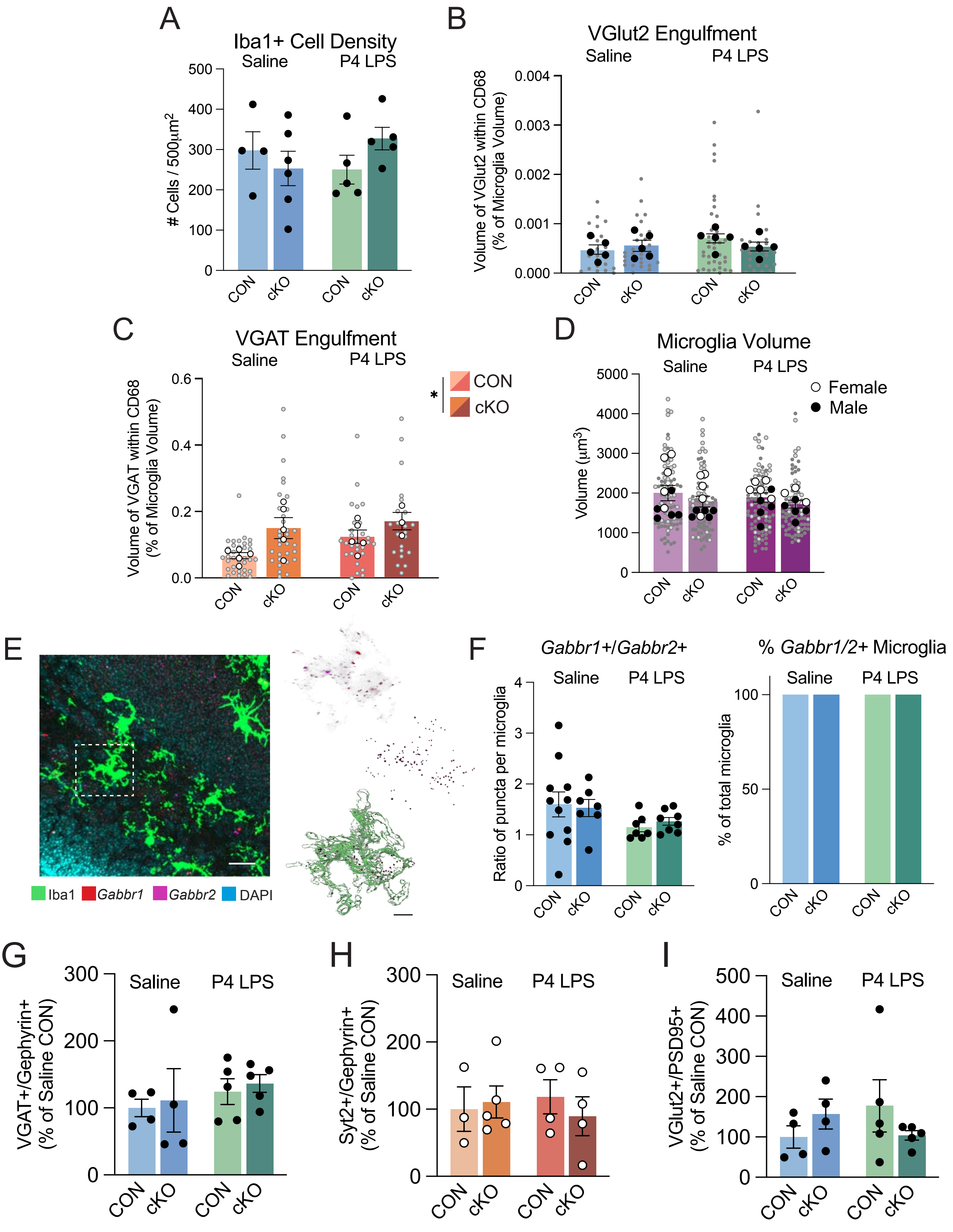

Supplementary Figure 2

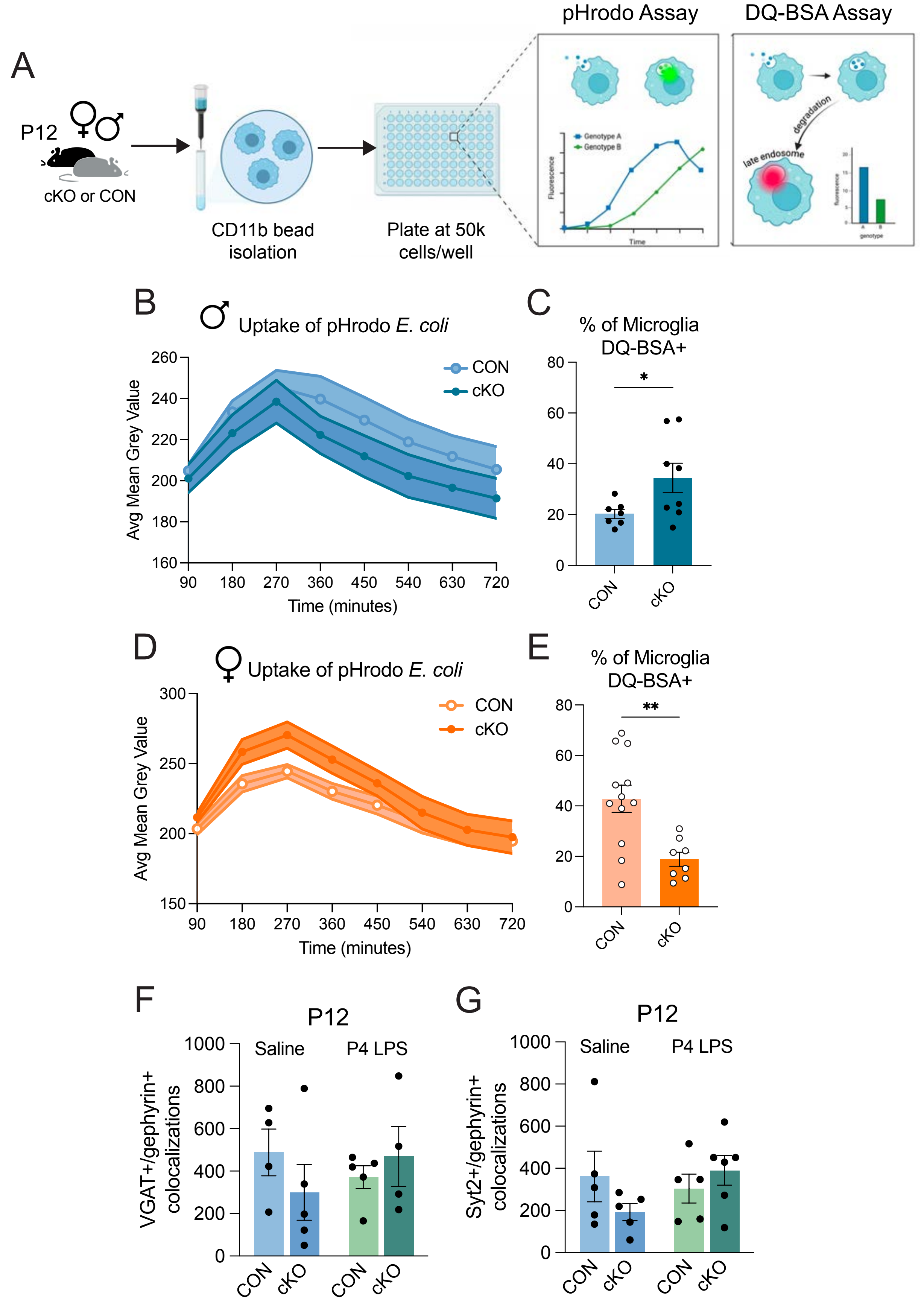

**Supplementary Figure 3**

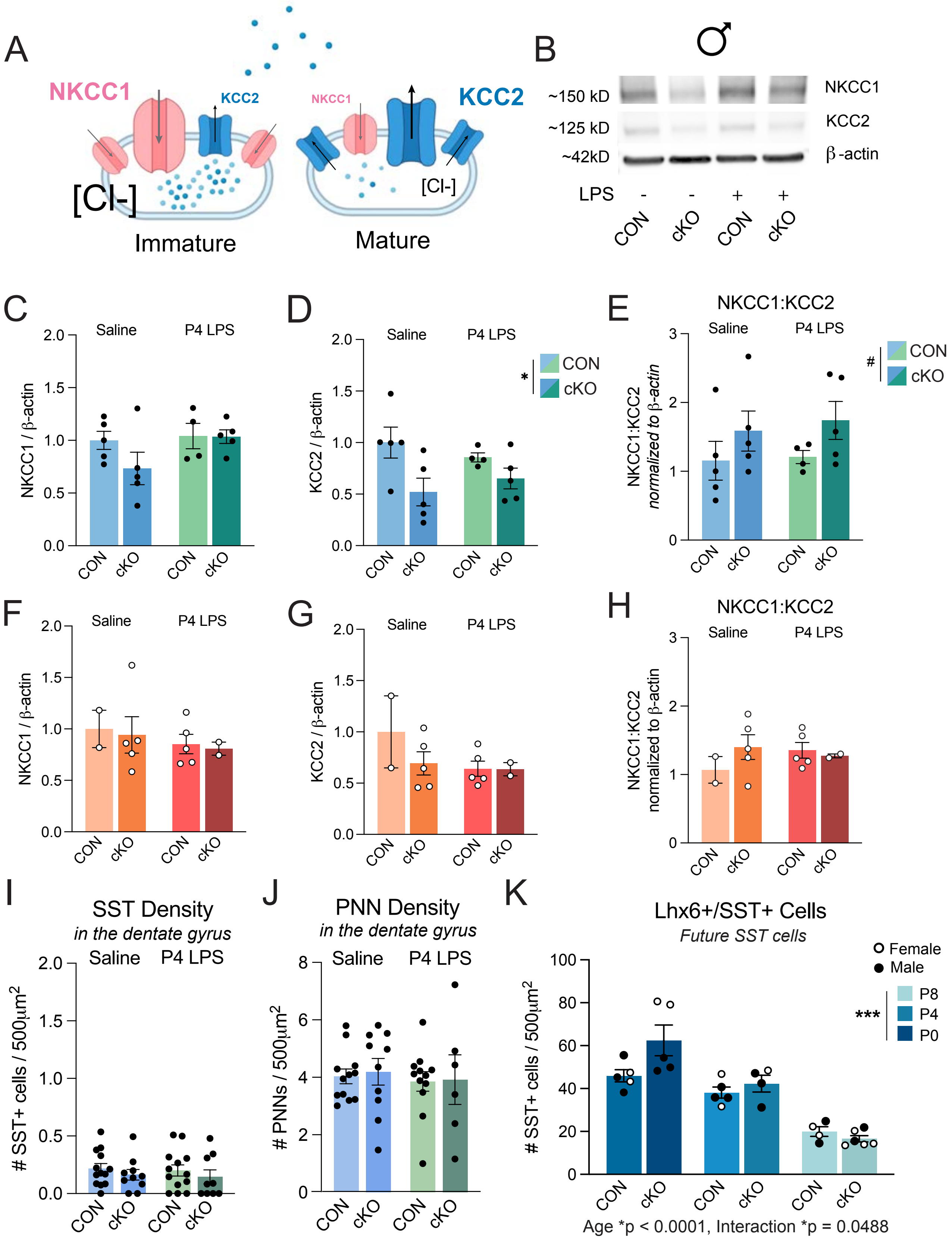

### Supplementary Figure 4

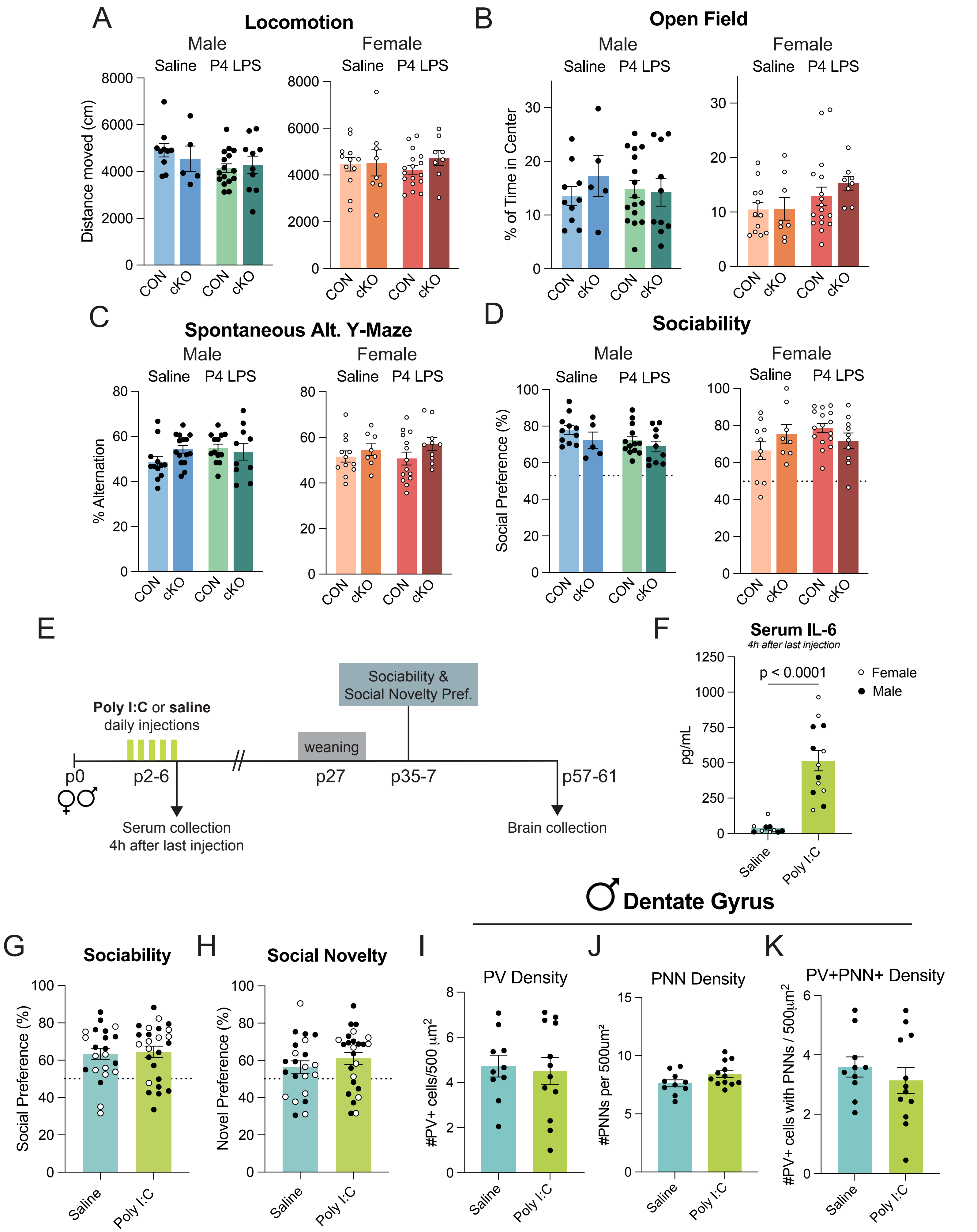

### Supplementary Figure 5

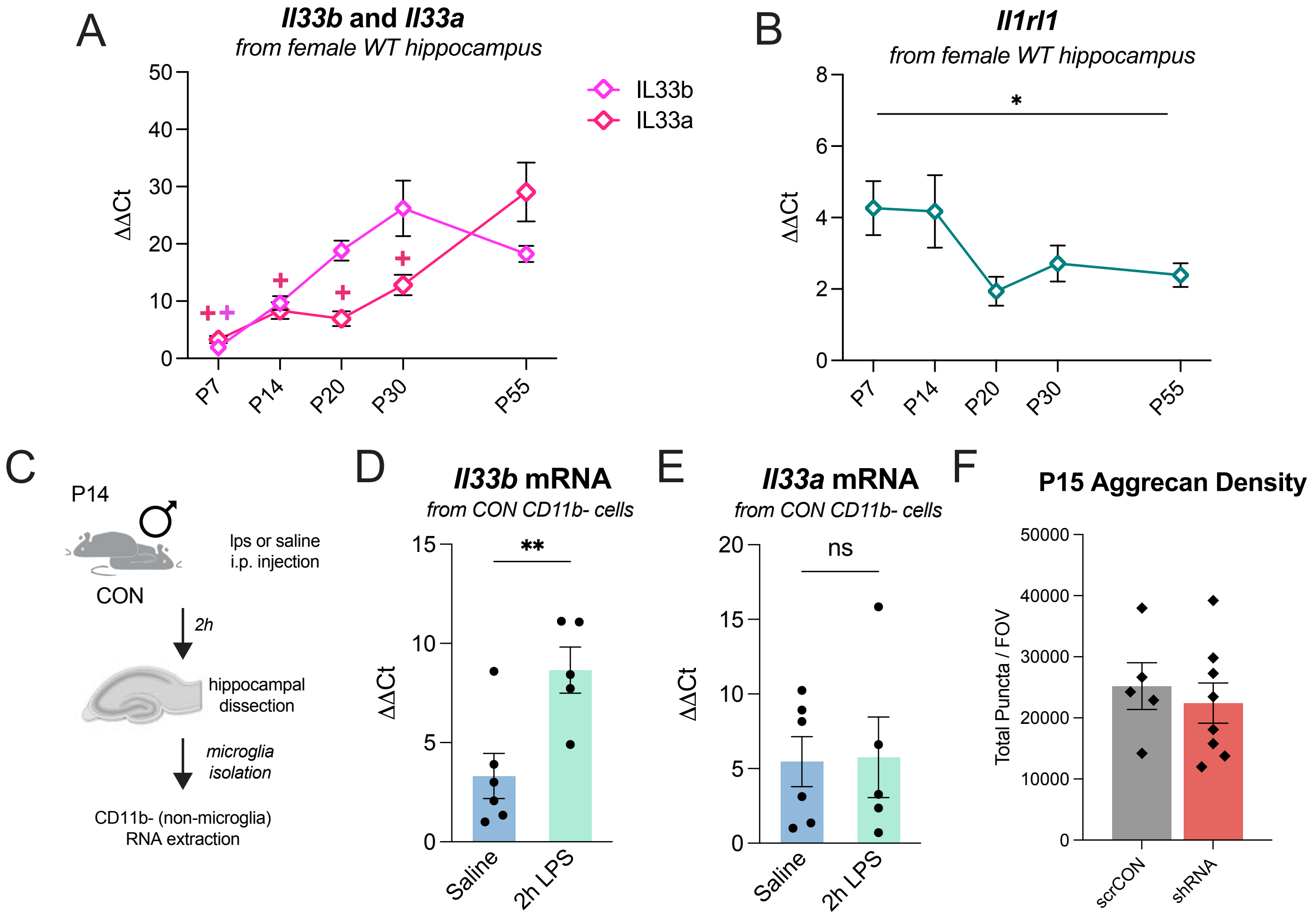
