## Supplemental Figure Legends 1-5 for "Microglial MyD88-dependent signaling influences extracellular matrix development and interneuron maturation in the hippocampus"

**Supplemental Figure 1. Characterization of male and female early postnatal MyD88-cKO microglia and synapses.**

**A.** Density of male microglia at P12 by Iba1+ somas, normalized to 500μm$^2$ area. Non-significant by 2-way ANOVA. **B.** Engulfment of excitatory VGlut2 presynaptic protein in male microglia lysosomes at P12, non-significant by 2-way ANOVA. **C.** Engulfment of inhibitory VGAT presynaptic protein within female microglia lysosomes at P12, significant main effect of genotype by 2-way ANOVA. **D.** Microglia 3D volume at P12 in males (closed circles) and females (open circles), non-significant by 2-way ANOVA. **E.** Left, representative image of P12 dentate gyrus, scale bar = 20μm. Right, image of *Gabbr1* (red) and *Gabbr2* (magenta) RNA signal within the inset microglia (top), "Spots" representation of RNA puncta (middle), and representative 3D reconstruction of microglia at P12 with Iba1 (green) protein (bottom), scale bar = 4μm. **F.** Quantification of *Gabbr1* and *Gabbr2* expression in dentate gyrus microglia, as ratio of *Gabbr1*:*Gabbr2* puncta per cell and percent of cells expressing both *Gabbr1* and *Gabbr2*, non-significant by 2-way ANOVA. **G.-I.** P18 quantification of synaptic density in males (**G., I.**) and females (**H.**) in the dentate gyrus. Non-significant by 2-way ANOVA. All error bars = SEM, all statistics run on individual animals (black or black-outlined circles), grey points represent individual microglia analyzed, except for *in situ* experiment (**F.**), where points represent individual microglia, from at least 3 animals/group.

**Supplemental Figure 2. P12 MyD88-cKO phagocytosis assays and inhibitory synapse density.**

**A.** Experimental design for phagocytosis assays with microglia isolated from P12 male and female mice. Microglia were plated to 50,000 cells per well, then treated with pHrodo *E. coli* or DQ-BSA and imaged in a live-imaging chamber. **B.** Male pHrodo uptake over time, non-significant by repeated measures 2-way ANOVA. **C.** Male percent of cells expressing DQ-BSA at 270 mins after treatment, significant unpaired t-test. **D.** Female pHrodo uptake over time, non-significant by repeated measures 2-way ANOVA. **E.** Female percent of cells expressing DQ-BSA at 270 mins after treatment, significant unpaired t-test. Each point = 1 animal, average of at least 3 wells/animal. **F.-G.** Inhibitory synapse density quantification in P12 males, non-significant by 2-way ANOVA. All error bars = SEM, all statistics run on individual animals.

**Supplemental Figure 3. Additional inhibitory system characterization in the MyD88-cKO hippocampus**

**A.** Schematic of chloride transporter concentrations over development, where increased KCC2 concentration and reduced NKCC1 concentration in maturity leads to normal intracellular chloride concentration, such that GABA signaling hyperpolarizes the cell. A high NKCC1 to KCC2 ratio is therefore considered immature. **B.** Representative western blots from males of NKCC1, KCC2, and β-actin for normalization. **C.** Male abundance of NKCC1, non-significant by 2-way ANOVA. **D.** Male abundance of KCC2, significant main effect of genotype by 2-way ANOVA. **E.** Ratio of male NKCC1:KCC2, trending (p=0.08) main effect of genotype by 2-way ANOVA. **F.-H.** Female quantifications of NKCC1 (**F.**), KCC2 (**G.**), and ratio (**H.**), all non-significant by 2-way ANOVA. **I.** Adult quantification of lowly-abundant somatostatin (SST) expressing interneurons in the male dentate gyrus, non-significant by 2-way ANOVA. **J.** Total number of perineuronal nets (PNNs) in the dentate gyrus of adult males, normalized to 500$\mu m^2$ area. Non-significant by 2-way ANOVA. **K.** Quantification of future SST expressing cells by *in situ* hybridization (*Lhx6*-/*Sst*+/DAPI+ cells) in the first postnatal week (P0-P8). Significant main effect of age and interaction (age x genotype) by 2-way ANOVA. All error bars = SEM, all statistics run on individual animals.

**Supplemental Figure 4. Additional adult behavioral characterization in MyD88-cKO males and females and early life poly I:C experiment behavior and brain endpoints**

**A.** Total distance moved in open field experiment over 5 minutes in males and females, non-significant by 2-way ANOVA. **B.** Time spent in center of open field apparatus in percent of 5-minute trial. Non-significant in males, trending main effect (p=0.05) of treatment in females by 2-way ANOVA. **C.** Percent correct alternations of total entries in Y-maze working memory paradigm, non-significant by 2-way ANOVA in males and females. **D.** Sociability assay quantified by percent social preference, calculated as time interacting with a social conspecific divided by total time spent interacting with social and non-social stimulus in the 3-chamber assay, *100. Non-significant in males and trending (p=0.05) genotype x treatment interaction in females by 2-way ANOVA. **E.** Experimental design for phenocopy early life poly I:C experiment. Male and female C57BL/6 wild-type mice were injected with poly I:C or saline daily from P2-P6, and social behavior was quantified prior to adult brain collection. Serum was collected in a subset of mice 4h after P6 poly I:C injection to verify immune response. **F.** Serum IL-6 expression by ELISA 4h after last injection, significant by unpaired t-test. **G.-H.** Sociability and social novelty preference, non-significant by unpaired t-test. **I.-K.** Male quantification of PV cell density (**I.**), PNN density (**J.**), and colabeled PV/PNNs (**K.**), normalized to 500μm$^2$ area. Non-significant by unpaired t-test.

**Supplemental Figure 5. Additional early life IL-33 characterization.**

**A.** Female *Il33b* and *Il33a* isoform expression across wild-type development. Significant main effects of age and isoform x age interaction effect by 2-way ANOVA. + indicates Bonferroni post-hoc significantly different than isoform-matched P55 adult expression. **B.** Female *Il1rl1* expression across wild-type development. Significant by one-way ANOVA, no significant post-hoc differences compared to P55 adult expression, n = 8 mice/age. **C.** Experimental design for non-microglia *Il33b* and *Il33a* expression following peripheral LPS injection. MyD88-CON male mice were injected with saline or 330ug/kg of LPS at P14, then microglia were isolated from hippocampal dissections. CD11b-negative (non-microglia) RNA was extracted and RT-qPCR was performed. **D.** LPS significantly increased *Il33b* expression by unpaired t-test. **E.** Acute peripheral LPS had no significant effect on *Il33a* expression. **F.** Density of total aggrecan puncta from the dentate gyrus (60x magnification) at P15 in mice previously treated with scrambled control virus or shRNA against *Il33* virus at P4, non-significant unpaired t-test. All error bars = SEM, all statistics run on individual animals.
